# AI-guided competitive docking for virtual screening and compound efficacy prediction

**DOI:** 10.1101/2025.10.28.685112

**Authors:** Manon Mirgaux, Valeria Barcelli, Adeline C.Y. Chua, Pablo Bifani, René Wintjens

## Abstract

Machine learning has transformed how we predict protein structures and interactions, but its full potential in drug discovery is only beginning to be realized. In this study, we demonstrate that advanced deep learning tools —such as AlphaFold3 and Boltz-1/2— not only predict protein-ligand interactions with high accuracy but can also separate active drug compounds from inactive ones. We present a straightforward strategy called *pairwise competitive docking*, which ranks drug candidates by directly comparing how well they bind to a protein’s target site. When applied to both bacterial and human enzymes, this method produced rankings that closely matched experimental results. We further show how this approach can guide the design of improved antibiotics and speed up the discovery of promising drug candidates from large chemical libraries. Overall, our findings highlight how machine learning can make structure-based drug design faster, more reliable, and more cost-effective.

## Introduction

Structure-based drug discovery has traditionally relied on virtual docking, but the advent of machine learning has dramatically transformed the field^1-3^. A breakthrough came with the development of denoising diffusion-based generative models^4^, leading to highly effective tools such as RoseTTAFold All-Atom^5^ and Alphafold3 (AF3)^6^. These tools can model protein-ligand interactions across a broad and unrestricted range of ligand types. Since then, several other similar tools have emerged, showing docking performances comparable to those of their precursors^7-10^.

A key challenge in machine learning-based docking predictions is that, regardless of the ligand type—whether a true binder or a non-interacting molecule—an interacting docking pose is typically generated between the target protein and the ligand. This raises two fundamental questions: (i) How can we distinguish true docking interactions from false positives? (ii) How can docking poses be ranked to reflect their expected binding affinities to a target protein?

We aimed to better harness the potential of machine learning-based methods by applying deep learning models such as AF3^6^ and Boltz-1/2^7, 10^ to well-characterized drug targets. We showed that these diffuse model-based programs can accurately reproduce docking poses and that, with appropriate criteria, it is feasible to distinguish true docking interactions from false positives. To support this, we developed a scoring procedure based on competitive docking to rank inhibitors targeting a specific binding site. We also demonstrated how the concept of competitive docking can be extended to virtual screening and drug design.

To showcase the versatility of our approach, we analyzed six systems: the tyrosine kinases TYK2 and CDK2, protein tyrosine phosphatase 1B, the cGMP-dependent 3’,5’-cyclic phosphodiesterase PDE2, four standard protein-ligand interaction benchmarks^10,11^, and the two human cyclooxygenases (COXs), also known as prostaglandin G/H synthases^12^. For a more complex test case involving multiple binding sites, we selected bacterial DNA gyrase because of its extensive characterization and the wealth of literature data available^13, 14^.

Our competitive docking approach produced rankings that strongly correlated with experimental inhibition data, underscoring the robustness of the method. In addition, we showed that simultaneous docking competition among a diverse set of compounds effectively identified the strongest binders, providing a powerful strategy for virtual screening hit discovery. Taken together, these findings open promising avenues for designing more effective inhibitors across a wide range of protein targets.

## Results

### Pose convergence can help identify real inhibitors

We used six human enzymes as benchmarks to see whether denoising diffusion models can separate true binders from false positives. After checking the available crystal structures for each benchmark protein (Table S1), we predicted binding poses for several inhibitors without experimental structures, and we also included 16 unrelated “off-target” compounds for each enzyme (Table S2). These off-target molecules normally bind to completely different proteins from those studies here.

We judged docking specificity using two measures: how close the ligands stayed to the binding site across the predicted models, and how consistent the ligand poses were with each other (docking convergence), measured as the average RMSD. Overall, real inhibitors bound in the pocket within about 3 Å and showed strong convergence, usually below 2 Å (Fig. 1). In contrast, off-target molecules were placed further away and with much higher variation.

**Fig. 1.**
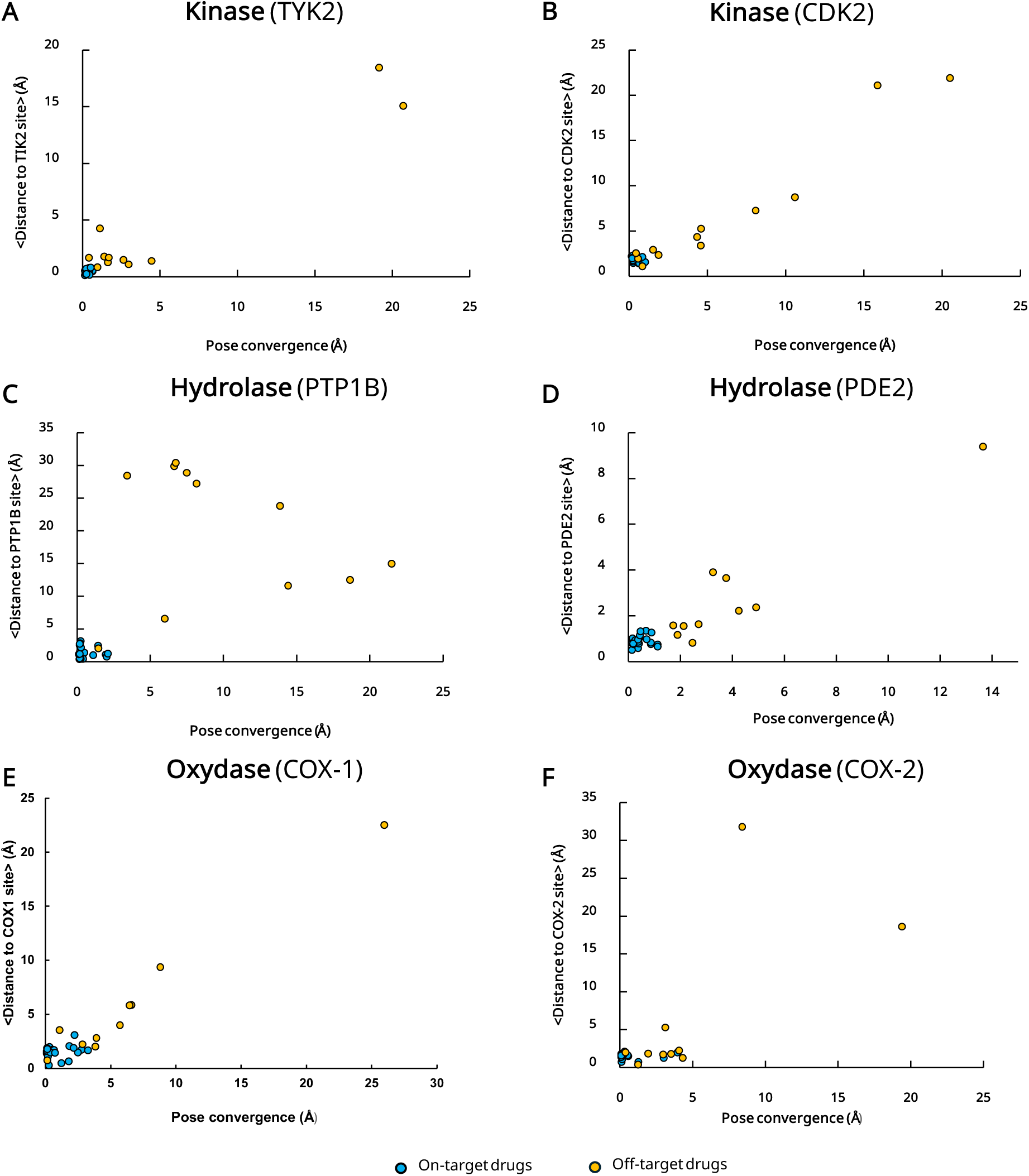
Docking specificity of AF3 across six protein benchmarks. Scatter plots show ligand pose convergence (RMSD) versus binding site distance for true on-target inhibitors (blue) and unrelated off-target compounds (orange). Results are presented for six enzymes: the kinases TYK2 (**A**) and CDK2 (**B**), the hydrolases PTP1B (**C**) and PDE2 (**D**), and the oxidases COX-1 (**E**) and COX-2 (**F**).

Specificity was especially strong for the TYK2 kinase and the two hydrolases, PTP1B and PDE2. In contrast, the weakest specificity was observed with CDK2 and COX-1, and to a lesser extent with COX-2, where some off-target molecules were still positioned near the binding site and showed low RMSD values. Similar results were obtained using the Boltz-2 AI model (Fig. S1).

### Docking specificity in a molecular system with multiple binding sites

We next asked whether AI-based docking can also help identify the true binding site of a ligand in proteins that have multiple binding pockets. To address this, we selected *Mycobacterium tuberculosis* (*Mtb*) DNA gyrase as a model system, given its complex binding landscape and extensive characterization^13, 14^. DNA gyrase contains several inhibitory sites that are targeted by chemically diverse compounds, making it an excellent case study for evaluating the potential of machine learning in drug discovery.

Fluoroquinolones (FQs) are the main class of inhibitors for this enzyme^15-17^, but inhibition can also be achieved by non-FQ compounds^18, 19^, including novel bacterial type IIA topoisomerase inhibitors (NBTIs)^20-22^ and thiophene-based molecules^23^ (here referred as DNA gyrase allosteric inhibitors).

After validating the available crystal structures (Table S1), we predicted binding poses for several FQs lacking experimental structures, as well as a set of non-FQ compounds. These included both ligands that bind DNA gyrase at sites distinct from the FQ pocket and anti-tuberculous agents known to act on entirely different protein targets (Table S2).

On average, FQs clustered close to their known binding site, with a mean distance of less than 2 Å in models generated (Fig. 2B and Table S3). A similar pattern was observed for non-FQ inhibitors, such as those in the spiropyrimidinetrione class, represented by QPT-1 (QPT) and zoliflodacin (ZLF) (Fig. 2C). These compounds also function through the FQ mechanism by binding at FQ site^18, 19^. NBTIs were consistently positioned within 2 Å of their known binding site, located approximately 10 Å from the FQ site and between the two scissile DNA bonds^22, 24^ (Figs. 2B and 2D). This result was observed using both AF3 and Boltz-1, but not Boltz-2 (Table S3).

**Fig. 2.**
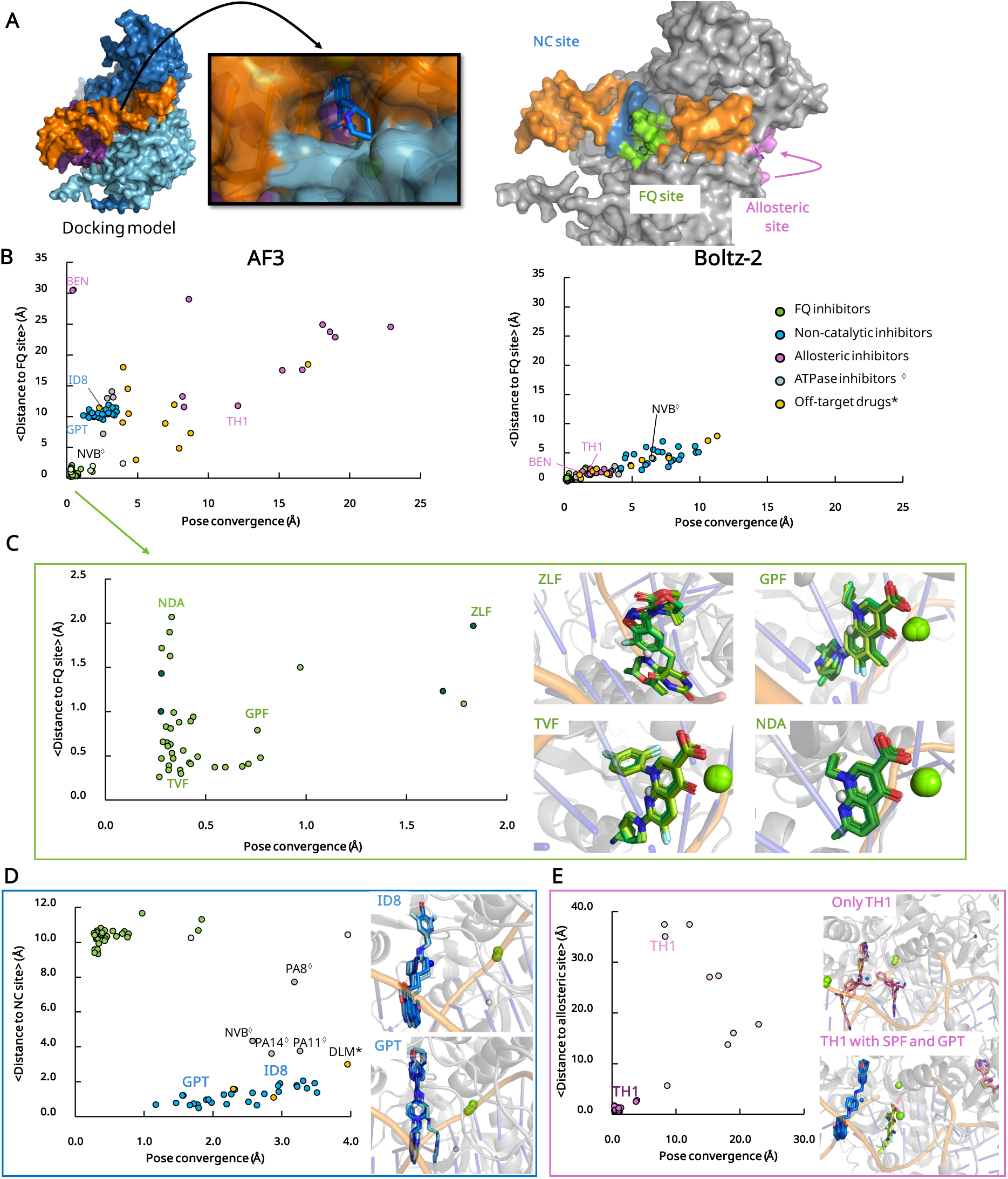
Docking specificity on *Mtb* DNA gyrase. (**A**) Molecular system used for DNA gyrase calculations (left) and closer view of the binding site localization (right). The three known binding sites in the studied molecular system are visualized. The gyrase system is shown as a gray molecular surface, DNA in orange, the FQ binding site in green, the non-catalytic (NC) site in blue, and the allosteric site in pink. (**B**) Clustering of DNA gyrase binders. AF3 (left) successfully distinguishes FQs (green), NC-site binders or NBTIs (blue), and allosteric inhibitors (pink) from unrelated compounds (orange and gray), based on docking pose convergence and proximity to the FQ site. Similar result was obtained with Boltz-1 (Table S3). In contrast, Boltz-2 (right) fails to achieve this separation, clustering all binders near the FQ site. Close-up of docking poses generated by AF3, for FQs (**C**), NBTIs (**D**), and allosteric inhibitors (**E**). A selected set of docked compounds is shown, including zoliflodacin (ZLF), grepafloxacin (GPF), trovafloxacin (TVF), nalidixic acid (NDA), AMK32b (ID8), gepotidacin (GPT), and thiophene 1 inhibitor (TH1). Each 3D image shows five representative AF3-predicted docking poses (i.e., the highest ipTM-scoring model for each seed) alongside center-of-mass of the reference FQ, MFX (gray sphere), superimposed on the reference crystal structure (PDB ID: 5BS8). Carbon atoms are color-coded as in (**B**), while oxygen and nitrogen atoms are shown in red and dark blue, respectively. The Mg^2+^ ions are represented as green spheres, the protein as a grey ribbon, and the DNA as orange phosphate backbone with green-blue base sticks. Both the protein and DNA are shown in transparency. Compounds marked with an asterisk (*) indicate drugs that do not target DNA gyrase or topoisomerase IV. Those marked with a rhombus (◊) are ATPase inhibitors, whose binding site is absent in the studied system.

In contrast, regardless of the diffusion models used, thiophene-class allosteric inhibitors were rarely docked at their expected binding site, located ∼30 Å from the FQ site (Fig. 2E)^23^. Instead, many of these inhibitors were mislocalized to the FQ or NC sites rather than their true allosteric site (Table S3). When docking was performed in the presence of gepotidacin (GPT) and sparfloxacin (SPF)—used to block the NC and FQ sites, respectively—the allosteric inhibitors remained confined to their proper binding site (Fig. 2E). Finally, although off-target compounds and gyrase ATPase inhibitors adopted plausible binding poses, they were, on average, significantly more distant from all three key binding sites of the molecular system under study (Fig. 2C).

Thus, the two criteria, pose convergence and proximity from the FQ site, effectively distinguished the three classes of gyrase inhibitors when suing AF3 and Boltz-1 diffusion models. The clustering pattern clearly separated FQs, NBTIs, and allosteric inhibitors—for these latter inhibitors, particularly when the other two binding sites were already occupied by their respective ligands—from unrelated compounds, which were more broadly dispersed across the clusters (Fig. 2).

### Competitive docking scoring and pairwise matrix

As shown in the previous section, diffusion-based models generate docking poses for any tested compounds, including those that do not specifically bind to the target protein. In the case of DNA gyrase, when two FQ molecules were docked simultaneously, only one occupied the catalytic FQ site, while the second often stacked against the DNA at the second catalytic site, which is only partially represented in the docking model used here. Furthermore, we previously showed that accuate docking of allosteric inhibitors on DNA gyrase can be achieved when both an FQ and an NBTI are included during docking inference (Fig. 2E). Building on these observations, we implemented a *pairwise competitive docking* approach to produce a scoring matrix, ranking compounds using a Competitive Docking Score (CDS) (Fig. 3).

**Fig. 3.**
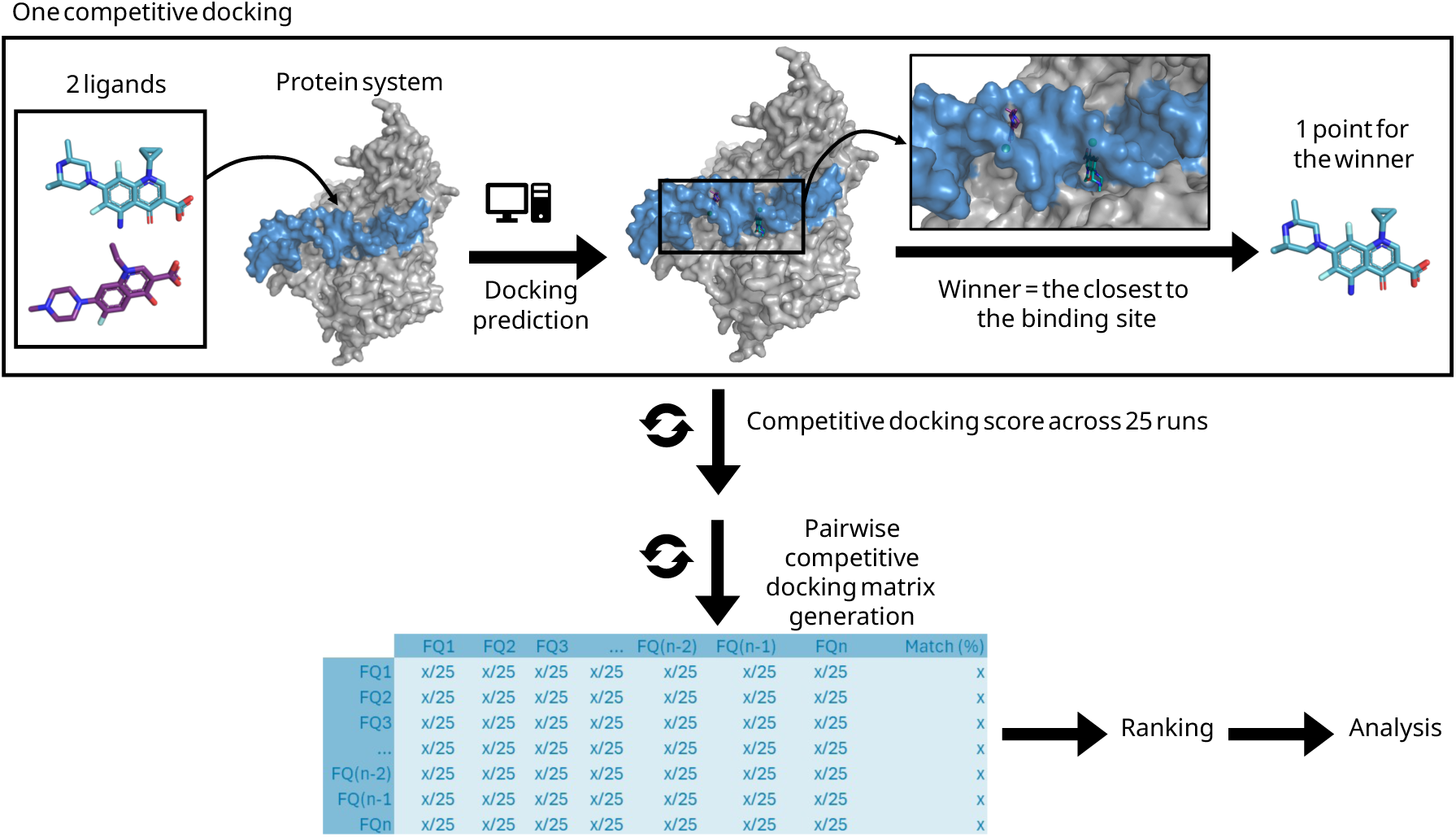
Scoring and ranking ligands with *pairwise competitive docking*. Diffusion-based machine learning predictions were performed using a protein model bound to two competing ligands. The ligand that successfully occupies the active site was considered the winner of each competitive docking run. A Competitive Docking Score (CDS) was calculated from at least 25 independent runs per ligand pair. These scores were compiled into a pairwise matrix to rank ligands based on their cumulative CDS. Expressed as a percentage, the final CDS reflects the win rate across all pairwise docking runs. As an example, Fig. S2 shows the CDS matrix obtained from competitive docking of 21 FQs on DNA gyrase using AF3.

### Competitive Docking Score correlation with inhibitor affinity

To evaluate the relevance of CDS rankings, we applied the method to benchmark proteins (Tables S1 and S2) and compared the results with experimental data reported in the literature^11, 20-28^ (Fig 4).

**Fig. 4.**
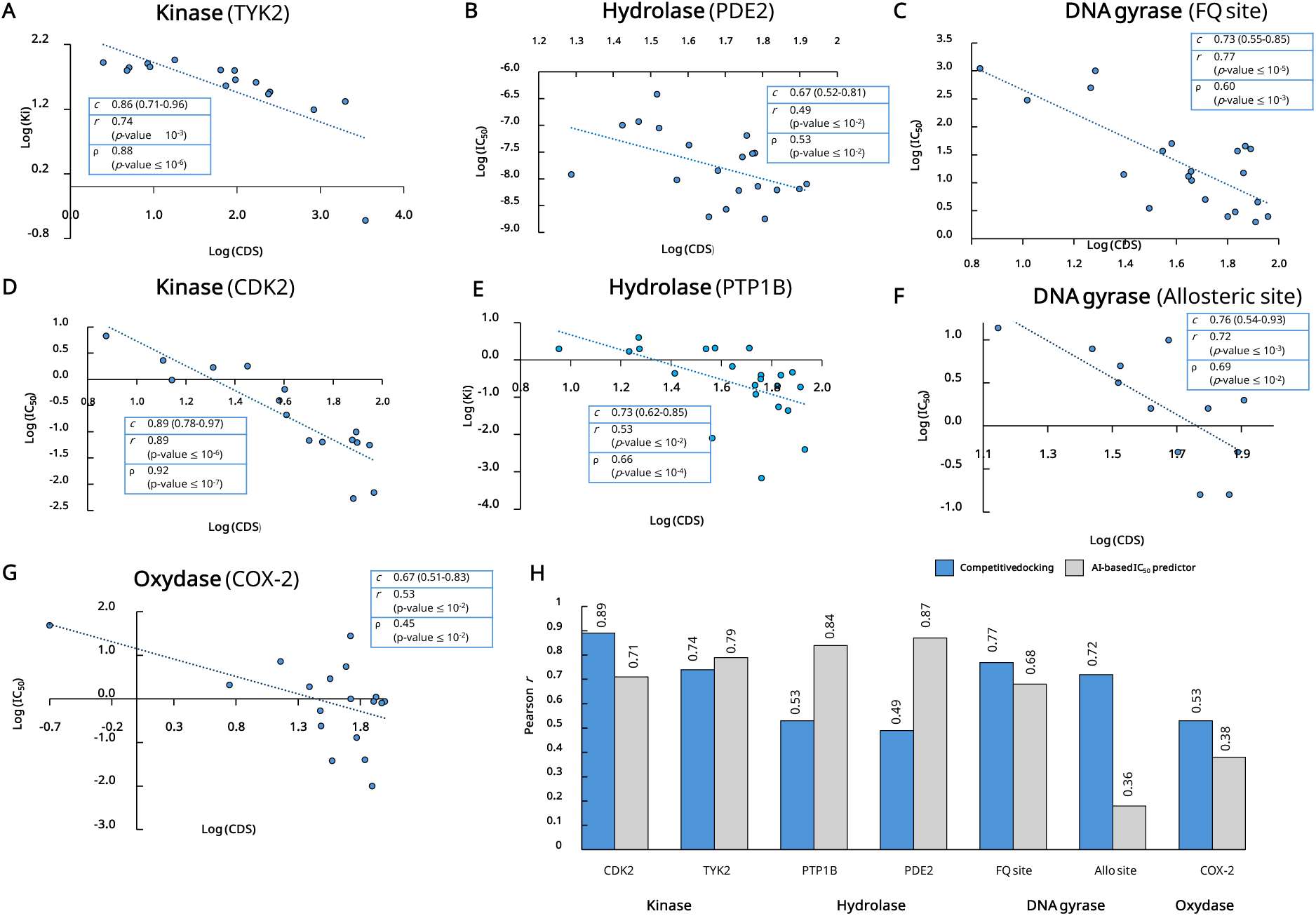
Correlation between competitive docking scores and experimental inhibitory activities across studied binding sites. **(A-G)** Scatter plots showing the relationship between the Competitive Docking Score (CDS) using AF3 and IC_50_ values for several systems studied. Each plot reports the Pearson correlation coefficient (*r*), Spearman’s rang correlation (ρ) with their corresponding *p*-values, and the rank concordance *c*-index with its 95% confidence intervale. An ordinary least squares regression line is added to illustrate the trend. **(H)** Comparison of AI-based strategies for predicting binding affinity. The competitive docking method was applied with AF3, while Boltz-2 was used as an AI-based IC_50_ predictor. Seven binding sites were evaluated, with performance summarized by the Pearson correlation coefficient (*r*).

Using AF3, the Pearson correlation coefficient (*r*) between CDS and experimental affinities ranged from very strong, with high statistical significance, for DNA gyrase FQs (*r* = 0.77, p < 0.00001), TYK2 inhibitors (*r* = 0.74, p < 0.001), and DNA gyrase allosteric inhibitors (*r* = 0.72, p < 0.001), to strong for PTP1B, COX-2, and PDE2 inhibitors (*r* = 0.53, 0.53, and 0.49, respectively), and moderate for NBTIs (*r* = 0.30) and COX-1 inhibitors (*r* = 0.24) (Fig. S3 and Table S4). Importantly, these latter two systems, which showed the weakest correlations, also displayed the lowest docking pose convergences (Figs. 1 and 2), suggesting that even without competition, inhibitors in these systems could not be reliably docked into the correct binding site.

It could be noted that *pairwise competitive docking* performed with Boltz-1/2 instead of AF3 generally produced weaker correlations with experimental IC_50_ values (Table S4), with no significant correlation observed for TYK2, PTP1B, and PDE2 systems.

We also found that the CDS rankings for COX-1 and COX-2 inhibitors were nearly identical (Pearson *r* = 0.96; Fig. S4), indicating that AF3 could not reliably distinguish between the two cyclooxygenase isoenzymes (Table S5). This limitation is likely due to the high degree of conservation between their binding sites (Fig. S5). To further examine this issue, we tested four major DNA gyrase variants known to confer resistance to FQs by increasing IC_50_ or MIC values by several-fold^25, 29^. In these cases, docking pose specificity remained largely unchanged, with only minor differences detected (Fig. S6). Similarly, the ranking of FQs with the *pairwise competitive docking* approach showed only slight shifts compared to wild-type results (Fig. S6). These findings suggest that substituting one or two amino acids in the catalytic site is not sufficient to alter AF3’s binding predictions.

### Competitive docking approach vs. direct AI-based affinity prediction

Recently, Boltz-2 was updated to include a module for predicting protein-ligand affinity. We compared the performance of this tool with our competitive docking approach by assessing their correlations with experimental data (Figs. 4G and 4H). Overall, both methods showed comparable performance across the eight tested systems, with only a few differences. For example, Boltz-2 performed better with the PTP1B and PDE2 systems, whereas competitive docking performed better for the DNA gyrase allosteric site, CDK2, and the COX-1/2 systems.

### DNA gyrase as a case study for evaluating CDS rankings

We applied the competitive docking method to 21 FQs within the *Mtb* DNA gyrase system and observed a strong correlation between CDS rankings and experimentally determined IC_50_ values using both AF3 and Boltz-1 (Figs. 4 and S5). Specifically, FQs with higher CDS values generally corresponded to more potent inhibitors, whereas those with lower CDS values were typically weaker (Figs. 5A and S6; Table S6). In contrast, Boltz-2 showed a weaker correlation, with a Pearson’s *r* of only 0.47 (Fig. S6).

**Fig. 5.**
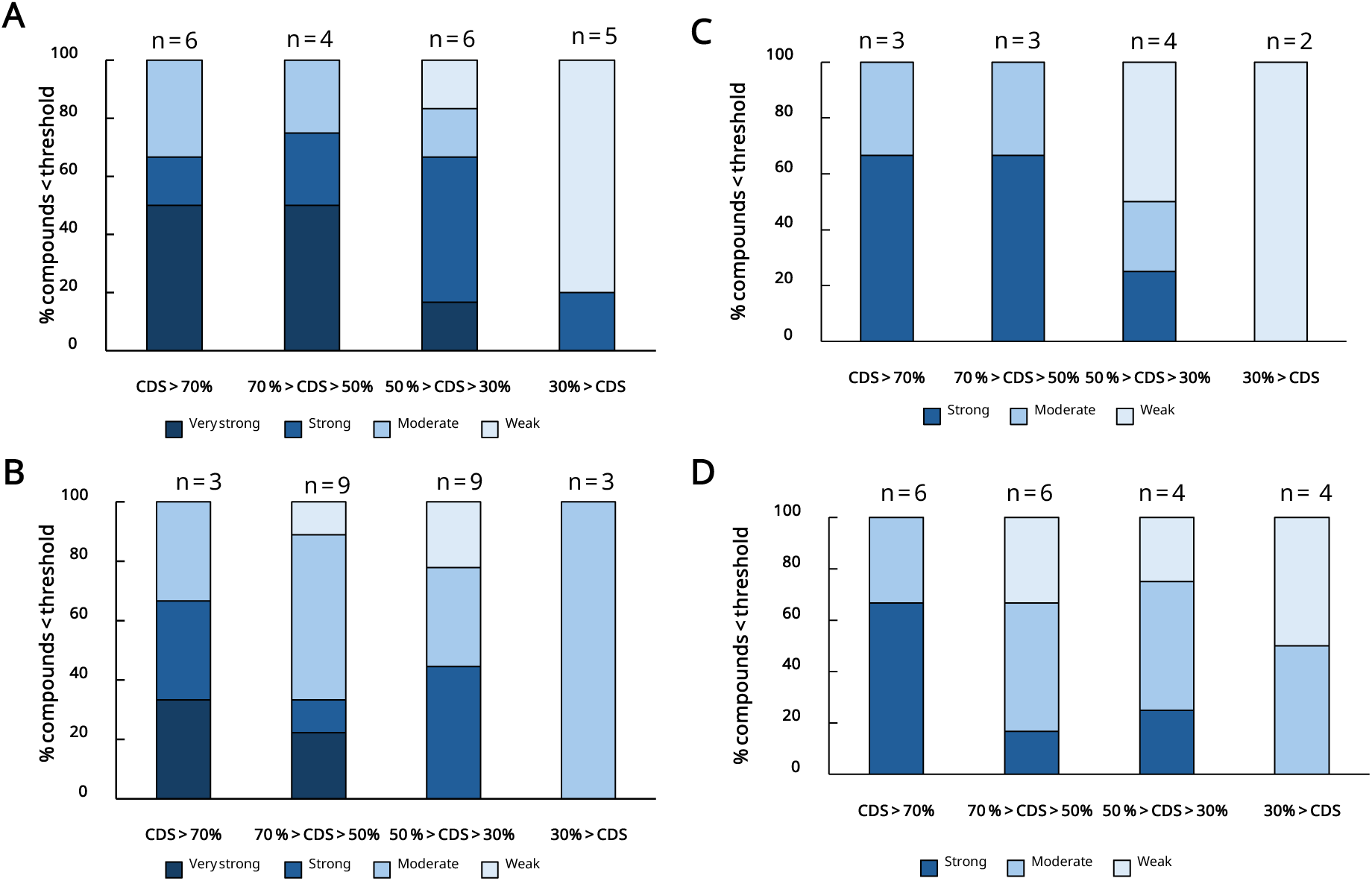
Distribution of DNA gyrase inhibitors across IC_50_ threshold categories, stratified by CDS ranking using AF3. **(A)** 21 FQs on *Mtb* DNA-gyrase, **(B)** allosteric inhibitors, **(C)** NBTIs, and **(D)** 20 FQs on *E. coli* DNA gyrase. The bar charts show the percentage of inhibitors within each IC_50_ threshold category, based on their CDS ranking. IC_50_ thresholds are provided in Table S4. ‘n’ indicates the number of inhibitors in each CDS category.

We next extended the approach to compounds targeting the two non-FQ binding sites in our gyrase model (Fig. 2A). For these analyses, we focused on systems with available experimental data—*E. coli* DNA gyrase for allosteric inhibitors (Table S7) and *S. aureus* DNA gyrase for NBTIs (Table S8). As with the FQs, stronger NBTIs and allosteric inhibitors consistently appeared in the high-CDS groups, while weaker inhibitors clustered in the low-CDS groups (Figs. 5B and 5C).

Because DNA gyrase is the primary FQ target in *E. coli*^30^, we also measured MIC_50_ values for the inhibition of *E. coli* growth by 22 FQs and compared them to their CDS rankings (Table S4 and Fig. S7A). Although the overall correlation was modest, a clear trend emerged: FQs with lower MIC_50_ values generally ranked higher in the CDS list (Fig. S7B). Notably, since FQs are known to have limited aqueous solubility^31^, excluding two FQs predicted to be poorly soluble significantly improved the correlation (Fig. 5D). Importantly, CDS rankings could not be further refined by considering the secondary FQ target in *E. coli*, as the rankings generated using *E. coli* topoisomerase IV were nearly identical to those based on DNA gyrase (*r* = 0.99) (Figs. S7D and S7E).

### *All-at-Once* docking strategy

To simplify the analysis and reduce computational cost, we evaluated AF3 docking performance by processing entire inhibitor classes in a single run, rather than relying on pairwise competitions. Using DNA gyrase as a model system, we tested three set of inhibitors with AF3: 21 FQs, 24 NBTIs, and 12 allosteric inhibitors. For the allosteric set, docking was performed in the presence of GPT and SPF to block the other two competing binding sites.

Overall, the *All-at-Once* strategy yielded less detailed results compared to the pairwise competitive approach (Table S9). While some of the top-ranked compounds based on CDS values showed strong occupancy within the target binding site, several potent inhibitors were not identified among leading competitors. Moreover, the number of distinct compounds effectively occupying the binding site was too limited for a reliable comparative analysis.

The *All-at-Once* strategy appears to be effective at identifying strong FQs from weaker FQs and non-FQ compounds. To evaluate its performance in a virtual screening context, we tested it on a compound library of 3,155 FDA-approved molecules, including 46 FQs, representing 1.5% of the total library. The library was randomly divided into 124 sets containing 25-26 compounds each, and *All-at-Once* docking was performed using AF3 (Fig. 6A).

**Fig. 6.**
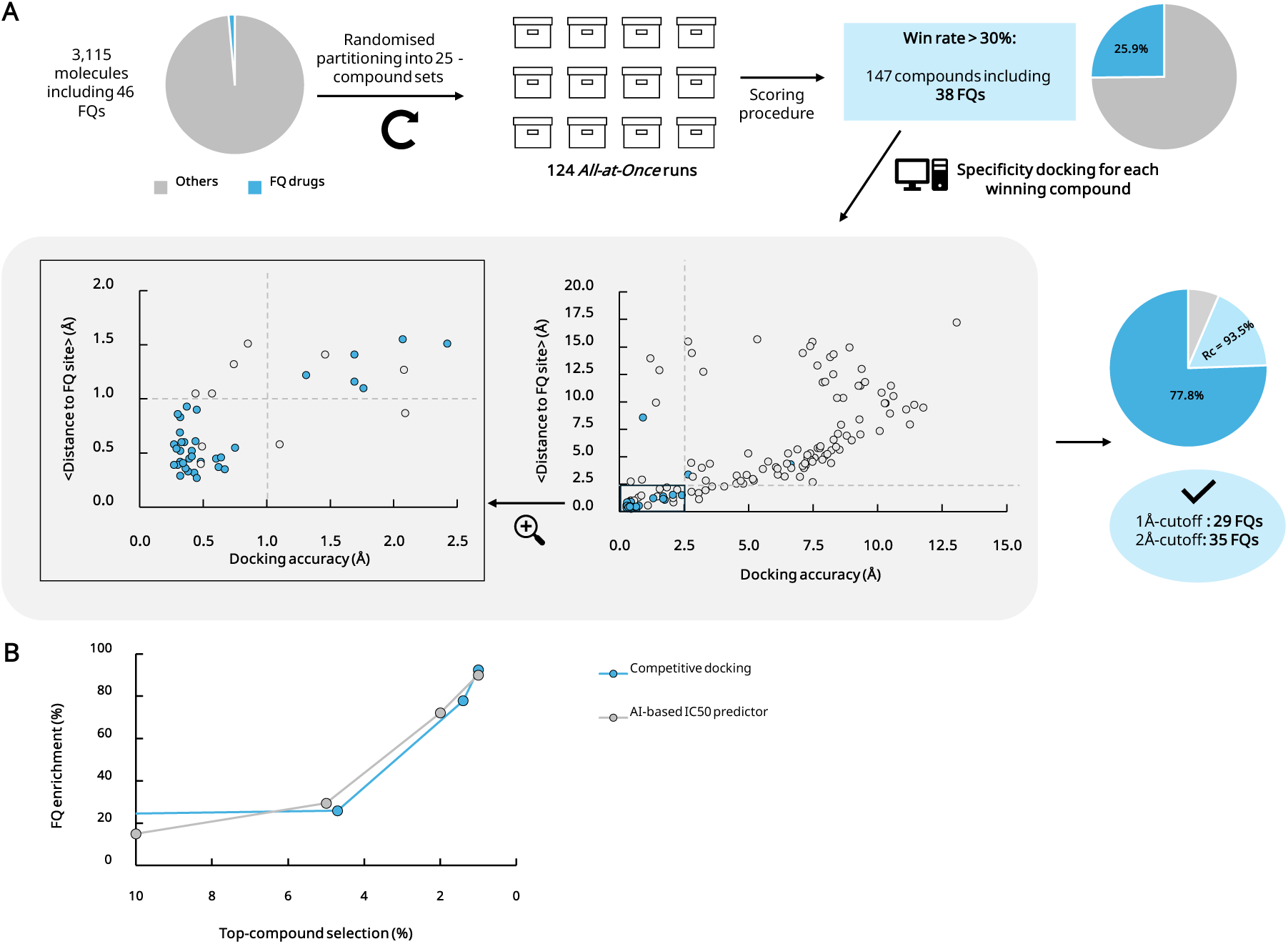
Finding effective FQ molecules with the *All-at-Once* strategy. (**A**) Virtual screening was performed on a library of 3,155 compounds, including 46 FQs (**B**) Percentage of FQs identified at different levels of the top-ranked compound list.

This screening identified 147 top-ranking compounds, including 38 FQs, corresponding to a 25.9% enrichment. The eight FQs not selected as winners are known to have low inhibitory activity against *Mtb* DNA gyrase^31^ and mostly belong to the first-generation FQs (Table S10). Applying an additional filter based on pose convergence (cutoff of 2.5 Å) and proximity to the FQ binding site (cutoff of 2.0 Å), as described in Fig. 2, increased FQ enrichment to 77.8% among the 45 remaining compounds. Tightening these thresholds to 1.0 Å further boosted enrichment to 93.5%, with an enrichment factor of 62. These results closely matched the performance obtained with Boltz-2 for hit identification across the 3,155-compound library (Fig. 6B).

### Applying competitive docking to design more potent FQs

Given that competitive docking can assist in identifying the most effective compounds for a specific target, we investigated how this approach could be employed to design more potent FQs. As proof of concept, and without exhaustive exploration, we selected a set of 414 compounds from several thousand automatically generated using the STONED algorithm^32^. This selection focused on the chemical space surrounding the five top-ranked FQs identified by AF3 (Table S10). Each *de novo* compound was then evaluated in competitive docking against STF, the highest-ranked FQ, using the *Mtb* DNA gyrase model system.

Thirty-one of these newly designed compounds occupied the FQ binding site in at least 70% of the 100 generated models, suggesting a stronger binding potential than STF. Their Tanimoto structural similarity to STF ranged from 0.25 to 0.88 (Fig. S8). Since none of these compounds are listed in the CAS chemical database—indicating they have likely never been synthesized—we further filtered them based on predicted ADME properties. Ultimately, only eight *de novo* compounds exhibited favorable drug-likeness characteristics, solubility, and chemical synthetic accessibility.

## Discussion

Artificial intelligence is rapidly reshaping *in silico* molecular docking workflows^33, 34^. Machine learning tools based on diffusion models have demonstrated remarkable accuracy in predicting protein-ligand interactions, often outperforming traditional docking programs^5-10^. In this study, we implemented a straightforward pairwise competitive docking approach to rank inhibitors of seven different enzymes, and nine total binding sites, using three pre-trained denoising diffusion-based models (Fig. 3).

Importantly, the effectiveness of our competitive docking approach depends on how the used deep learning models adhere to physicochemical laws—particularly in managing atomic clashes within the generated structures. As AF3 includes a clash penalty in its model ranking, such failures are less frequent in AF3-generated models compared to those produced Boltz-1 or Boltz-2. The steering versions of both latter models effectively removed atom clashes in the generated models, but without significant improvement in competitive docking performance (Table S4). Hence, overall, AF3-based competitive docking outperformed the steering Boltz models across all cases (Fig. S1). Other diffusion-based models such as RoseTTAFold All-Atoms^4^, Chai^8^, or Protenix^9^, or NeuralPlexer^35^ should be evaluated to assess their potential in competitive docking scenarios.

Across the binding sites analyzed, the competitive docking approach performed on par with the recently released affinity prediction module in Boltz-2, both in hit-to-lead inhibitor ranking mode (Fig. 4G) and in screening hit identification mode (Fig. 6B). To enable a more robust comparison between the two methods, additional systems will need to be evaluated. Notably, in cases where both approaches performed poorly—such as the DNA gyrase NC site, COX-1 and COX-2—the competitive docking method still outperformed Boltz-2. This underscores a key limitation of current AI-based tools: they are not universally reliable across all targets. Certain systems may consistently underperform for reasons that remain unclear, which we plan to investigate further in future studies. Thus, our competitive docking strategy provides a valuable alternative when direct AI-based affinity predictors fail or when dealing with particularly challenging systems.

Our scoring procedure, based on competitive docking within an identified binding site, offers a simple and interpretable alternative to more complex tools that depend on external scoring functions^36^. Although the concept of competitive docking is not entirely new, previous applications have typically focused either on placing multiple ligands side by side within large binding pockets^37^ or on fragment-based drug design^38, 39^. To the best of our knowledge, using a machine learning-based tool to simulate virtual competition between ligands for the same binding site is a novel strategy. Furthermore, our method provides a streamlined and accessible implementation leveraging machine learning models, with the introduction of novel evaluation criteria based on binding site occupancy. These criteria can be adapted to the specific system under investigation, and we plan to explore this flexibility further in future studies.

We also explored a multi-ligand competition strategy—referred to as *All-at-Once*—to identify top binders from a mixed set of weak and non-binding compounds. When applied to a library of 3,115 compounds, this approach effectively selected strong FQs, while weaker ones were outcompeted during docking (Fig. 6A). Final enrichment of FQs reached 93.5% after applying a filter based on docking convergence derived from non-competitive predictions. The *All at-once* strategy offers substantial time savings compared to relying solely on non-competitive docking. Rather than evaluating all 3,115 compounds individually, only the 147 candidates selected by the *All-at-Once* approach needed further analysis for docking convergence. This efficiency gain becomes even more significant when applying the two-step process to larger compound libraries. Finally, as proof of concept, we applied our pairwise competitive docking method to design improved inhibitors targeting the FQ binding site. Several *de novo* compounds were selected with eight predicted to have favorable drug-like properties, including ADME profiles, solubility, and synthetic accessibility. These chemically novel candidates merit synthesis and experimental validation in future studies.

## Methods

### Biological systems investigated

This study primarily examines the interactions between various seven proteins and various compounds: two human kinases (TYK2, CDK2), two human hydrolase (PTP1B and PDE2), two human oxidases (COX-1 and COX-2) and the bacterial DNA gyrase.

For all the models, the sequences used were based on corresponding crystal structures deposited on the PDB. The human tyrosine kinase (TYK2), the human cyclin-dependent kinase 2 (CDK2), human phosphodiesterase 2A (PDE2) and the human tyrosine-protein phosphatase 1B (PTP1B) were considered as monomer using the complex crystal structures 4GIH^40^, 1H1Q^41^, 6C7E^42^ and 2QBS^43^ as reference template for the binding site, respectively.

In the COX model system, we analyzed a single monomer, which included a protoporphyrin IX moiety with a cobalt cation, based on the crystal structure of COX-2 bound to mefenamic acid (MFN) (PDB ID: 5IKR^44^). Finally,

Concerning the DNA gyrase/DNA complex, to optimize computational efficiency, the system was simplified to include:

- The N-terminal catalytic core of GyrA (∼500 residues).
- The C-terminal TOPRIM domain of GyrB (∼250 residues).
- A 95-residue GyrA fragment completing the active site with the catalytic residue Y129, which forms a transient O-(5’-phospho-DNA)-tyrosine covalent intermediate^45^.
- A 24-mer double-stranded DNA molecule.
- Two Mg^2+^ cations.

The DNA sequences were derived from the crystal structure of the *Mtb* DNA gyrase/DNA complex bound to MFX (PDB ID: 5BS8^46^). Additionally, four major *Mtb* GyrA variants associated with FQ resistance were incorporated into the docking predictions: G88A, A90V, D94G, and the double mutant A90V+D94G^25, 29^. DNA gyrase and Topoisomerase IV (Topo IV) from *Escherichia coli* and *Staphylococcus aureus* were considered using the same double-stranded DNA sequence, corresponding protein fragments, and Mg^2+^ ions as in the *Mtb* gyrase model.

### Molecular compounds investigated

A total of 189 compounds were examined covering a range of inhibitors for TYK2, CDK2, PTP1B, PDE2, COXs and DNA gyrase:

- TYK2, CDK2, PDE2 and PTP1B inhibitors (16, 16, 21 and 22 compounds, respectively): these inhibitors were taken from the standard protein-ligand benchmark dataset^11^.
- NSAIDs (22 compounds): these were used to assess the COX system, including diclofenac (DCF), ibuprofen (IBU), and paracetamol (PCT).
- FQs (34 compounds): five of these have known *Mtb* DNA-gyrase/DNA/FQ crystal structure.
- Non-catalytic inhibitors (NBTIs) (26 compounds): these bind midway between the two FQ binding sites. Two compounds have an *Mtb* DNA gyrase co-crystal structure (PDB ID: 9FOY^47^).
- Allosteric inhibitors (12 compounds): these bind to GyrA at an allosteric site, stabilizing the DNA-cleavage complex similarly to FQs^23^.
- ATPase inhibitors (4 compounds): targeting the ATPase domains of topoisomerases, including novobiocin (NVB).
- Non-FQ inhibitors targeting the FQ site (5 compounds): including zoliflodacin (ZLF), etoposide (ETP), and QPT-1 (QPT), the first discovered spiropyrimidinetrione^18^.
- Anti-tuberculosis drugs (9 compounds): first-line and second-line TB drugs, including bedaquiline (BDQ), delamanid (DLM), ethambutol (ETB), ethionamide (ETH), isoniazid (INH), linezolid (LNZ), pyrazinamide (PZA), rifampicin (RIF), and streptomycin (STR).
- Two negative-control compounds were included: the antipsychotic aripiprazole (ARI) and the anticancer agent bosutinib (BOS).

The smile name of the compounds investigated are reported in Table S2. These compounds were used in their protonated forms at pH 7.0 predicted with MarvinSketch.

In addition, a library of 3,155 compounds was used for *in silico* screening. This library comprises FDA-approved and pharmacopeia drugs from the commercial collection offered by TargetMol Chemicals Inc. (Massachusetts, USA, Catalog No. L1010). Twenty-two FQs, along with the two negative-control compounds, were purchased from TargetMol Chemicals for experimental MIC testing against *E. coli*.

Finally, *de novo* compounds were automatically generated by exploring the chemical space surrounding the five top-ranked FQs using the STONED algorithm^32^. Several hundred compounds were produced, and those with a Tanimoto similarity coefficient ≥ 0.75— indicating close structural similarity to the top FQs—were selected for further investigation in this study.

The ADME properties of these compounds were predicted using the SwissADME web server^48^. The SMILES representations and structural similarity of the *de novo* compounds to 5 top-ranked FQs are provided in Table S12. Structural similarity was evaluated using the Tanimoto coefficient calculated from ECFP4 fingerprints generated with RDKit (https://www.rdkit.org).

### Machine learning calculations

Boltz-1 and Boltz-2 were installed locally from its official GitHub repository (https://github.com/jwohlwend/bolt). The steered versions of both programs, i.e. that apply physics-based potentials to enhance the physical plausibility of the structures, were also considered^10^. The AF3 source code was retrieved from GitHub on November 14, 2024 (https://github.com/google-deepmind/alphafold3), and model parameters were obtained directly from Google. Public sequence and structure databases were downloaded from AlphaFold’s storage. Computational resources included: GPU-based model inference using NVIDIA RTX 4080 (16GB), RTX 3090 (24GB) or A40 (48GB), and CPU-based sequence and template searches using an 11^th^ Gen Intel 8-core CPU with 128 GB RAM.

To optimize computational time, multiple sequence alignments were pre-generated for each molecular system, and non-protein components (DNA, ions, and compounds) were incorporated before model inference. At least five randomly selected seed numbers were used, generating a total of a minimum of 25 models per model inference.

### Pose evaluation metrics

Docking performance was evaluated by superimposing predicted proteins onto reference crystal structure using PyMol (super command, aligning a-carbons), then ligand heavy-atom root-mean-square-deviation (RMSD) values between docked ligands were computed using DockRMSD^49^:

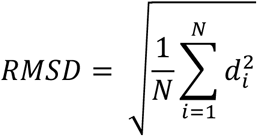

where N is the number of heavy atoms in the ligand, and *d*_*i*_ is the Euclidean distance in angstroms (Å) between the ith pair of corresponding atoms.

Three performance metrics are considered:

- Accuracy (Å): mean RMSD between predicted and experimental poses.
- Precision (Å): convergence of docking results, measured by mean pairwise RMSD across generated models.
- per chain ipTM score for the top-ranking prediction: the per chain interface predicted template modeling (ipTM) score measured the accuracy of the predicted relative positions of the two chains considered within the complex model. ipTM is on a 0-1 scale, with values higher than 0.8 representing high-quality predictions, while values below 0.6 suggest a failed prediction^6^.

### Pairwise competitive docking

To rank compounds based on their binding affinity to a specific binding site, we employed a pairwise competitive docking approach. This method involves running docking predictions with two compounds simultaneously in a single inference step.

Following protein alignment with reference structure, we measured the center of mass distances between each docked compound and the reference complex. The compound positioned closest to the reference ligand’s binding site was considered to adopt the correct binding pose and was designated as the winner for that site. A 5 Å cutoff was applied, meaning that if the winning compound’s distance exceeded this threshold, the competition was deemed inconclusive. Additionally, docking simulations where the two competing compounds exhibited steric clashes (defined as any interatomic distance below 1.54 Å) were also classified as inconclusive and excluded from further analysis.

For each pairwise comparison, we generated 25 independent docking models using randomized seed values, evaluating outcomes across these 25 generated poses. The results were compiled into a pairwise competition matrix, which was then used to rank compounds based on their Competitive Docking Score (CDS) (Fig. 3). The CDS reflects the percentage of wins for each compound within the matrix.

For TYK2, CDK2, PDE2 and PTP1B, the binding site reference points were calculated with the ligand in the reference structure. For COX system, the binding site reference point was calculated as the average center of mass of ligand found in PDB IDs 5IKR and 5KIR^44^. In the DNA gyrase system, we used the center of mass of MFX from the reference structure as the FQ site reference point. Concerning the two additional binding sites for non-FQ inhibitors were also considered: for the NC site, the reference point was derived from AMK32b’s center of mass in PDB ID 9FOY^47^, superimposed onto the DNA gyrase reference structure; and for allosteric site, the reference point was based on the benzoisoxazole **3** ligand from PDD ID 6QX1^23^.

### *E. coli* growth inhibition screening

*Escherichia coli* strain ATCC 25922 was cultured in LB broth at 37 °C with shaking until mid-log phase (OD₆₀₀ = 0.5–0.7), then diluted to a final assay OD₆₀₀ of 0.01. Test compounds from TargetMol Chemicals Inc. (Massachusetts, USA)—including 22 FQs and two negative-control drugs—were prepared as 10 mM or 1 mM stock solutions in DMSO or water, as appropriate. These were serially diluted in 384-well “mother” plates to 50× the final assay concentrations and transferred to assay plates containing either the bacterial inoculum or LB broth for control wells.

Final DMSO concentrations did not exceed 2% (v/v), which did not affect bacterial growth. Plates were incubated overnight at 37 °C in sealed, humidified containers. Growth inhibition was assessed by measuring OD₆₀₀ relative to untreated controls. MIC_50_ values were determined using four-parameter nonlinear regression (GraphPad Prism v10.0) and are reported as mean ± standard deviation from at least three independent biological replicates.

### *All-at-Once* calculations

In addition to pairwise competitions, we evaluated AF3’s docking performance when multiple compounds were processed simultaneously in a single inference run. In this *All-at-Once* approach, 100 structural models were generated per run. The compound most frequently observed within the FQ binding site was considered the “winner”, and a win rate was calculated for each input compound.

As in the pairwise competition, steric clashes were assessed using the same distance cutoff (1.54 Å). Docking attempts involving a clash were considered inconclusive and excluded from further analysis. Due to the high frequency of atomic overlaps in models generated by Boltz-1, the *All-at-Once* strategy was implemented exclusively using AF3.

For the virtual screening of the 3,155-compound library including 46 FQs, we randomly divided the dataset into 109 sets of 25 compounds and 15 sets of 26 compounds, yielding 124 *All-at-Once* runs. A win-rate cutoff of 30% was applied to identify top-performing competitive compounds for the FQ site. This threshold was selected under the assumption that the presence of four FQs in a single set was highly improbable (estimated probability: 0.04% based on the hypergeometric distribution), while the probability of three FQs occurring in the same set was approximately 0.5%.

Boltz-2x was also applied in virtual screening mode of *Mtb* DNA gyrase using the 3,155-compound library, with compounds ranked according to the binding likelihood predicted by Boltz-2.

### Statistical evaluation

The CDSs and rankings obtained were correlated with the experimental inhibition values. Correlations were assessed against available IC_50_ and MIC values. For DNA gyrase and Topo IV systems, IC_50_ represents the drug concentration required to inhibit DNA supercoiling by 50%. MIC values correspond to the drug concentration that reduces bacterial growth to 1% or less compared to the drug-free control culture. IC_50_ for COX-1 and COX-2 were collected from the literature based on human whole blood assays.

To assess statistical correlations, we used three metrics: Pearson’s correlation coefficient (*r*), which measures the strength of a linear relationship between two numerical variables and is sensitive to the magnitude of differences, Spearman’s rank correlation (ρ), which evaluates monotonic trends based on rank order rather than actual values, and the concordance index (*c*-index or *c*), which quantifies the proportion of all comparable pairs for which predictions and experimental outcomes are ranked in the same direction. This latter metric is particularly relevant in our study, where the correct ranking of compounds is more important than accurately predicting their exact inhibitory values.

The *c*-index was defined using the following formula:

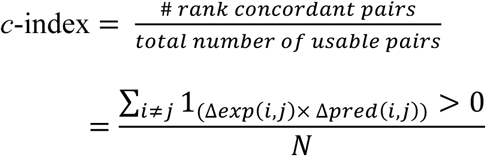

where i and j are the ranks of two observations, Δexp(i, j) represents the difference in experimental ranks, Δpred(i, j) represents the difference in predicted docking ranks, *N* is the total number of comparable pairs, and a pair (i, j) contributes a value of 1 if the product Δ*exp*(*i*, *j*) x Δ*pred*(*i*, *j*) > 0 (i.e., the predictions are concordant with the experimental outcomes), and 0 otherwise. A 95% confidence interval for the c-index was computed using the bootstrap method with 1,000 resamples of the dataset with replacement.

The statistical significance of the *r* and ρ correlations was determined using a t-test, following the formula:

Pearson t-test:

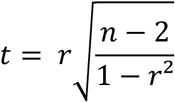

where *r* is the Pearson’s correlation coefficient and n is the number of data points.

Spearman t-test:

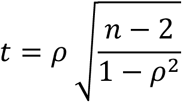

where *n* is the number of data points and ρ is the Spearman’s rank correlation.

The t-statistic follows a t-distribution with n-2 degrees of freedom, allowing us to calculate a two-tailed *p*-value.

## Data availability

All data necessary to support the conclusions in this study are provided in the main text and/or the Supplementary Materials. Key datasets and source code are available upon request from the authors and can also be accessed a https://doi.org/10.5281/zenodo.15241961.

## Acknowledgments

Computations for this work were carried out both locally and, in part, on the Hercules2 cluster of the CÉCI high-performance computing infrastructure. We extend our special thanks to Frédéric Wautelet for his assistance with the installation of AF3 on this platform. The authors are also grateful to Dr Alex Wohlkönig and Prof. Anna-Maria Marini for their valuable discussions and critical review of the initial manuscript. MM is a research Fellow and RW is a Research Associate, both funded by the Belgian Fund for Scientific Research (FNRS), whose support is gratefully acknowledged. PB is funded by grants NRF-CG2025-CG02-IG2-001001 and supported by the Lee Kong Chian School of Medicine - Ministry of Education Start-Up Grant.

## Author contributions

M.M., P.B., and R.W. conceived the study. R.W. supervised the project. M.M., V.B., A.C.Y.C. and R.W. carried out the experiments and acquired data. All authors analyzed data and wrote the article.

## Competing interests

The authors declare no competing interests.

## Materials & Correspondence

Correspondence and requests for materials should be addressed to

## Notes

### Competing Interest Statement

The authors have declared no competing interest.

