## Supplementary data file for "AI-guided competitive docking for virtual screening and compound efficacy prediction"

Manon Mirgaux *et. al.*

**This PDF file includes:**

Tables S1 to S11

Fig. S1 to S8

References

**Table S1. AF3-predicted poses for selected crystallographic structures of the studied proteins.** *Precision* is reported as the mean pairwise RMSD of ligands across 25 generated models, while the *Accuracy* corresponds to the mean RMSD between the AF3-predicted poses and the crystallographic reference pose. The per-chain ipTM metric was calculated specifically at the protein-ligand interface. *

| Protein | Ligand name | Ligand ID | PDB ID | RSR | RSCC | *Precision (± SD)* | *Accuracy (± SD)* | *Per chain ipTM* |
|---|---|---|---|---|---|---|---|---|
| CDK2 | NU6094 | L1H1Q | 1H1Q | 0.105 | 0.895 | 0.26 (± 0.10) | 0.xx (± 0.xx) | 0.97 (± 0.01) |
| TYK2 | xx | xx | 4GIH | 0.120 | 0.920 | 0.xx (± 0.xx) | 0.xx (± 0.xx) | 0.xx (± 0.xx) |
| PTP1B | xx | xx | 2QBS | 0.080 | 0.980 | 0.xx (± 0.xx) | 0.xx (± 0.xx) | 0.xx (± 0.xx) |
| PDE2 | xx | xx | 6C7E | 0.102 | 0.960 | 0.xx (± 0.xx) | 0.xx (± 0.xx) | 0.xx (± 0.xx) |
| COX-2 | Meclofenamic acid | MCN | 5IKQ | 0.068 | 0.961 | 0.55 (± 0.92) | 0.56 (± 0.67) | 0.96 (± 0.01) |
| | Mefenamic acid | MFN | 5IKR | 0.070 | 0.938 | 0.08 (± 0.03) | 0.45 (± 0.03) | 0.96 (± 0.01) |
| | Tolfenamic acid | TFN | 5IKT | 0.063 | 0.968 | 0.52 (± 1.46) | 0.69 (± 0.99) | 0.95 (± 0.03) |
| | Flufenamic acid | FFN | 5IKV | 0.068 | 0.956 | 0.17 (± 0.12) | 0.38 (± 0.06) | 0.97 (± 0.01) |
| | Vioxx/Rofecoxib | RFC | 5KIR | 0.262 | 0.858 | 0.08 (± 0.03) | 0.84 (± 0.01) | 0.96 (± 0.01) |
| DNA-gyrase | 8-methyl-moxifloxacin | 8MX | 5BTN<br>5BTL | 0.343<br>0.273 | 0.889<br>0.860 | 0.54 (± 0.38) | 0.76 (± 0.27) | 0.91 (± 0.01) |
| | ID8 | A1ID8 | 9FOY | 0.225 | 0.893 | 3.59 (± 2.33) | 3.67 (± 0.38) | 0.71 (± 0.03) |
| | ID9 | A1ID9 | 9FOY | 0.222 | 0.901 | 4.02 (± 2.49) | 3.86 (± 0.45) | 0.69 (± 0.03) |
| | Ciprofloxacin | CPF | 5BTC | 0.322 | 0.910 | 0.30 (± 0.09) | 0.57 (± 0.07) | 0.91 (± 0.01) |
| | Gatifloxacin | GFN | 5BTD | 0.233 | 0.938 | 0.42 (± 0.42) | 0.58 (± 0.25) | 0.90 (± 0.01) |
| | Levofloxacin | LFX | 5BTG<br>5BTI | 0.222<br>0.249 | 0.920<br>0.931 | 0.28 (± 0.09) | 0.56 (± 0.08) | 0.90 (± 0.01) |
| | Moxifloxacin | MFX | 5BTA<br>5BS8 | 0.291<br>0.180 | 0.888<br>0.945 | 0.71 (± 0.42) | 0.78 (± 0.33) | 0.91 (± 0.01) |

*Accuracy* and *Precision* values are in Å. Electron density fit metrics, the real-space R factor (RSR) and the real-space correlation coefficient (RSCC), were averaged over the observed ligand in the crystallographic unit. These metrics were retrieved from the RCSB Protein Data Bank. For clarity, the ligand in 9FOY is designed ID8 (S stereoisomer) and ID9 (R stereoisomer).

**Table S2. SMILES representations of the molecules studied for each protein.** The dataset includes inhibitors of CDK2 kinase (16), TYK2 kinase (16), tyrosine phosphatase 1B (PTP1B, 22), phosphodiesterase 2A (PDE2, 21), cyclooxygenases COX-1 and COX-2 (22), and DNA gyrase across its three binding sites (76). In addition, 16 off-target compounds were included. In total, 189 compounds are listed with their corresponding SMILES strings.

**< TableS2-Smiles.xlsx>**

**Table S3. Docking specificity of AF3 and Boltz-1/2 for the DNA gyrase system.** For each diffusion-based model (AF3, Boltz-1, Boltz-2), pose Accuracy and Precision are reported as defined in Table S1. Distances to each of the three binding sites were calculated. Successful docking is expressed as the percentage of poses located within 5 Å of the corresponding target-inhibitor binding site.

**< TableS3-AF3 and Blotz-1 docking specificity.xlsx>**

**Table S4. Experimental inhibition data of the studied inhibitors in relation to their Competitive Docking Score (CDS) for the corresponding target protein.** CDS values from pairwise competitive docking with AF3, Boltz-1, or Boltz-2 are reported alongside experimental inhibitory values ($IC_{50}$ or $K_i$). Nine systems were analyzed: CDK2, TYK2, PTP1B, PDE2, COX-1, COX-2, and the three binding sites of DNA gyrase. The Pearson correlation coefficient (*r*) and Spearman's rank correlation coefficient (ρ) are also provided.

**< TableS4-AF3 and Boltz1-2 CDS score.xlsx>**

**Table S5. Relationship between Competitive Docking Score (CDS) from AF3 and the inhibitory potency of COX-1 and COX-2 inhibitors.** Experimental inhibition data[17] were obtained using a whole blood assay and compared with their CDS values. COX-1 inhibitors are classified as strong ($IC_{50}$: 0.01–1.0 µM), moderate ($IC_{50}$: 1.0–30.0 µM), or weak ($IC_{50}$: >30.0 µM). For COX-2, the classification is strong ($IC_{50}$: 0.01–1.0 µM), moderate ($IC_{50}$: 1.0–5.0 µM), or weak ($IC_{50}$: >5.0 µM).

| Ligand name | Code | COX-1 | | | COX-2 | | |
|---|---|---|---|---|---|---|---|
| | | Range | $IC_{50}$ (µM) | CDS (%) | Range | $IC_{50}$ (µM) | CDS (%) |
| Celecoxib | CLX | Moderate | 6.7 | 95.9 | Strong | 0.87 | 99.8 |
| Compound 13 | CO1 | Weak | 100 | 33.4 | Strong | 0.24 | 30.5 |
| Compound 25 | CO2 | Moderate | 5.8 | 92.7 | Strong | 0.80 | 94.1 |
| Diclofenac | DCF | Strong | 0.075 | 57.4 | Strong | 0.038 | 37.5 |
| Dup-697 | DUP | Strong | 1.0 | 57.1 | Strong | 0.01 | 78.8 |
| Etoricoxib | ERX | Weak | 116 | 82.6 | Moderate | 1.1 | 84.7 |
| Fluorbiprofen | FBF | Strong | 0.075 | 48.0 | Weak | 5.5 | 48.7 |
| Ibuprofen | IBU | Moderate | 7.6 | 10.2 | Weak | 7.2 | 14.4 |
| Indomethacin | IDT | Strong | 0.013 | 43.5 | Moderate | 1.0 | 52.9 |
| Ketoprofen | KPF | Strong | 0.047 | 37.3 | Moderate | 2.9 | 36.2 |
| Lumiracoxib | LRX | Weak | 67 | 38.5 | Strong | 0.13 | 59.1 |
| Meloxicam | MLX | Moderate | 5.7 | 16.4 | Moderate | 2.1 | 5.6 |
| Naproxen | NPX | Moderate | 9.3 | 66.0 | Weak | 28 | 52.7 |
| Nimesulide | NMS | Moderate | 10 | 58.2 | Moderate | 1.9 | 24.7 |
| Paracetamol | PCT | Weak | >100 | 0.4 | Weak | 49 | 0.2 |
| Rofecoxib | RFC | Moderate | 19 | 24.3 | Strong | 0.53 | 30.1 |
| SC-58125 | SC5 | Moderate | 13 | 53.4 | Strong | 0.04 | 68.9 |
| Valdecoxib | VDX | Moderate | 26 | 84.6 | Strong | 0.87 | 81.2 |

**Table S6. Experimental inhibition data of *Mtb* DNA gyrase and corresponding Competitive Docking Scores (CDS) for 21 FQs.** Experimental values were taken from Aubry et al.[16], while CDS values were obtained using AF3, Boltz-1, or Boltz-2 diffusion-based models. Inhibitors are classified as very strong ($IC_{50}$: 2-5 µg/ml), strong ($IC_{50}$: 5-20 µg/ml), moderate ($IC_{50}$: 20-50 µg/ml), or weak ($IC_{50}$: > 50 µg/ml). Solubility class was predicted using the SILICOS-IT software.

| Ligand | Inhibitor range | $IC_{50}$ (µg/ml) | MIC (µg/ml) | 21 FQ CDS with AF3 (%) | 21 FQ CDS with Boltz-1 | 21 FQ CDS with Boltz-2 | Solubility class |
|---|---|---|---|---|---|---|---|
| SPF | Very | 2 | 0.25 | 81.2 | 56.4 | 48.0 | Soluble |
| STF | Very | 2.5 | 0.25 | 90.8 | 58.6 | 48.0 | Moderate |
| CNF | Very | 2.5 | 0.5 | 63.2 | 59.0 | 69.0 | Soluble |
| GFN | Very | 3 | 0.12 | 67.6 | 54.1 | 51.0 | Soluble |
| CPF | Very | 3.5 | 0.5 | 31.2 | 50.2 | 80.0 | Soluble |
| MFX | Very | 4.5 | 0.5 | 82.6 | 50.5 | 43.0 | Soluble |
| LFX | Very | 5 | 0.5 | 51.6 | 56.8 | 27.0 | Soluble |
| GMF | Strong | 11 | 4 | 45.6 | 54.4 | 62.0 | Soluble |
| GRX | Strong | 13 | 2 | 44.4 | 58.7 | 55.0 | Poor |
| NRF | Strong | 14 | 4 | 24.8 | 39.4 | 56.0 | Soluble |
| TVF | Strong | 15 | 16 | 72.8 | 69.8 | 54.0 | Moderate |
| GPF | Strong | 16 | 1 | 45.4 | 60.1 | 55.0 | Moderate |
| PFX | Moderate | 37 | 8 | 35.2 | 45.7 | 40.0 | Soluble |
| TSF | Moderate | 37 | 16 | 69.0 | 70.4 | 43.0 | Moderate |
| TMF | Moderate | 40 | 4 | 77.6 | 65.7 | 48.0 | Poor |
| FRX | Moderate | 45 | 6.25 | 74.0 | 54.5 | 64.0 | Soluble |
| ENX | Moderate | 50 | 8 | 38.2 | 47.6 | 63.0 | Soluble |
| OXO | Weak | 300 | 64 | 10.4 | 21.6 | 37.0 | Soluble |
| FLQ | Weak | 500 | 64 | 18.4 | 36.9 | 40.0 | Soluble |
| PPM | Weak | 1000 | 128 | 19.2 | 26.5 | 42.0 | Soluble |
| NDA | Weak | 1100 | 128 | 1100 | 13.2 | 25.0 | Soluble |

**Table S7. Experimental inhibition data for *E. coli* DNA gyrase ($IC_{50}$) and *E. coli* growth inhibition (MIC), compared with Competitive Docking Scores (CDS) from AF3 for 12 allosteric inhibitors[13].** Inhibitors are classified as strong ($IC_{50}$: 0.16-1.0 μM), moderate ($IC_{50}$: 1.0-5.0 μM), or weak ($IC_{50}$: > 5.0 μM).

| Drug | Inhibitor category | $IC_{50}$ (μM) | MIC (μg/ml) | CDS with GPT and STF (ionized form) (%) | CDS with GPT and STF (non-ionized form) (%) |
|---|---|---|---|---|---|
| C12 | Strong | 0.16 | 0.25 | 72.9 | 88.6 |
| BEN | Strong | 0.16 | 0.12 | 59.2 | 42.4 |
| C11 | Strong | 0.5 | 0.25 | 77.4 | 55.0 |
| C09 | Strong | 0.5 | 1 | 50.5 | 13.4 |
| C04 | Moderate | 1.6 | 2 | 62.6 | 61.1 |
| C10 | Moderate | 1.6 | 8 | 41.6 | 36.2 |
| TH2 | Moderate | 2 | 16 | 80.9 | 61.6 |
| C06 | Moderate | 3.2 | 4 | 33.0 | 48.4 |
| C05 | Moderate | 5 | 2 | 33.5 | 79.8 |
| C07 | Weak | 7.9 | 4 | 27.4 | 71.0 |
| C08 | Weak | 10 | 2 | 47.4 | 37.6 |
| TH1 | Weak | 13.8 | 4 | 14.0 | 5.2 |

**Table S8. Experimental inhibition data for *S. aureus* DNA gyrase compared with Competitive Docking Scores (CDS) from AF3 for 24 NBTIs[6-11].** MIC data are provided for *S. aureus* (ATCC 29213) strains. NBTIs are classified as very strong ($IC_{50}$: 0.004–0.02 μM), strong ($IC_{50}$: 0.01–0.25 μM), moderate ($IC_{50}$: 0.25–2.0 μM), or weak ($IC_{50}$ > 2.0 μM). When $IC_{50}$ data were unavailable, classification was assigned based on MIC values, with corresponding ranges indicated in parentheses.

| Drug | Range | $IC_{50}$ (μM) | MIC (μg/ml) | CDS (%) |
|---|---|---|---|---|
| NC9 | Very strong | 0.007 | 0.031 | 42.0 |
| N10 | Very strong | 0.011 | 0.0078 | 65.9 |
| 423 | Very strong | 0.014 | / | 78.2 |
| 237 | (Strong) | / | 0.06 | 72.5 |
| AMK | Strong | 0.035 | 0.125 | 39.2 |
| NC7 | Strong | 0.067 | 0.125 | 49.2 |
| N16 | Strong | 0.16 | 0.125 | 40.0 |
| N21 | Strong | 0.16 | 0.25 | 60.1 |
| N15 | Strong | 0.24 | 0.5 | 53.9 |
| 587 | (Moderate) | / | 0.125 | 78.2 |
| N13 | Moderate | 0.33 | 0.25 | 65.2 |
| N20 | Moderate | 0.33 | 0.5 | 40.9 |
| N22 | Moderate | 0.34 | 0.5 | 34.2 |
| N19 | Moderate | 0.35 | 1 | 67.1 |
| GPT | Moderate | 0.37 | 0.125 | 82.0 |
| N12 | Moderate | 0.51 | 2 | 67.8 |
| N11 | Moderate | 0.55 | 1 | 33.6 |
| NC8 | Moderate | 1.02 | 4 | 12.4 |
| N17 | Moderate | 1.1 | 2 | 22.0 |
| NC6 | Moderate | 1.6 | 1 | 57.0 |
| N18 | Weak | 1.82 | 2 | 27.6 |
| NC1 | Weak | 4.39 | 4 | 37.5 |
| NC5 | Weak | 100 | >128 | 31.4 |
| N14 | Weak | 100 | >128 | 52.7 |

**Table S9. Competitive success rates using the *All-at-Once* docking strategy with AF3.** The table reports the percentage of binding site occupancy for each compound tested. The "/" symbol indicates compounds that were not included in a given competing set. Each AF3 inference run was performed with 20 seeds, generating 100 models. The compound achieving the highest success rate is shown in bold.

| **Site** | **Win rate (%)** | | | | | | | | | | | | | | |
|---|---|---|---|---|---|---|---|---|---|---|---|---|---|---|---|
| **FQ on *Mtb* DNA gyrase** | **STF** | **SPF** | **FRX** | **TMF** | **MFX** | **TVF** | **LMF** | **TSF** | **8MX** | **GFN** | **CPF** | **LFX** | **OXO** | **GPF** | **NDA** |
| 21 FQs | 13.5 | 0.0 | 0.0 | 39.2 | 41.9 | 0.0 | / | 4.1 | 0.0 | 0.0 | 1.4 | 0.0 | 0.0 | 0.0 | 0.0 |
| Screening on 27 ligands[1] | **73.7** | / | / | / | 25.0 | / | / | / | / | / | / | 0.0 | 0.0 | 1.3 | 0.0 |
| **NC on *S. aureus* DNA gyrase** | **GPT** | **423** | **587** | **237** | **AMK** | **NC1** | **NC5** | **NC6** | **NC7** | **NC9** | **N10** | **N12** | **N13** | **N14** | **N22** |
| 24 NBTIs | 9.8 | 65.6 | 1.6 | 1.6 | 8.2 | 0.0 | 0.0 | 6.6 | 0.0 | 1.6 | 0.0 | 1.6 | 1.6 | 0.0 | 1.6 |
| Screening on 23 ligands[2] | 6.5 | **75.5** | / | / | / | 0.0 | / | / | / | 4.3 | / | 13.7 | / | / | / |
| **Allosteric on DNA gyrase** | **BEN** | **C04** | **C05** | **C06** | **C07** | **C08** | **C09** | **C10** | **C11** | **C12** | **TH1** | **TH2** | **PA11** | **PA14** | **BDQ** |
| All allosteric on *Mtb* | 0.0 | 0.0 | 2.0 | 0.0 | 0.0 | 0.0 | 0.0 | 0.0 | 0.0 | 0.0 | 0.0 | 98.2 | / | / | / |
| Screening on *E. coli* with 25 ligands[3] | 0.0 | / | / | / | 28.9 | / | / | / | / | 26.3 | / | 9.2 | 2.6 | 1.3 | **31.6** |
| **COX** | **CLX** | **SC5** | **FBF** | **IBU** | **PCT** | **DCF** | **IDT** | **C01** | **C02** | **DUP** | **ERX** | **KPF** | **LRX** | **MLX** | **NMS** |
| 22 NSAIDs on COX-1 | 83.7 | 10.2 | 0.0 | 0.0 | 0.0 | 1.0 | 0.0 | 0.0 | 0.0 | 1.0 | 3.1 | 0.0 | 1.0 | 0.0 | 0.0 |
| 22 NSAIDs on COX-2 | 65.6 | 22.9 | 0.0 | 0.0 | 0.0 | 0.0 | 0.0 | 2.1 | 0.0 | 0.0 | 7.3 | 0.0 | 2.1 | 0.0 | 0.0 |
| Screening on 32 ligands with COX-1[4] | **100** | / | 0.0 | 0.0 | 0.0 | 0.0 | 0.0 | / | / | / | / | / | / | / | / |
| Screening on 32 ligands with COX-2[5] | **64.0** | 36.0 | 0.0 | 0.0 | 0.0 | 0.0 | / | / | / | / | / | / | / | / | / |

[1] Screening is composed of 9 FQs (STF, MFX, GPF, NDA, OXO, CNX, FLQ, PPM, LFX) and 18 off-target molecules (CLX, SC5, FBF, IBU, PCT, NVB, PA8, P11, P14, ETB, INH, PZA, ETH, STR, DLM, BDQ, LNZ, RIF).

[2] Screening is composed of 5 NBTIs (GPT, 423, NC1, NC9, N12) and 18 off-target molecules (ETB, INH, PZA, ETH, NVB, ETP, STR, QPT, IP3, IP1, DLM, ZLF, PA8, P11, P14, BDQ, LNZ, RIF).

[3] Screening is composed of 5 allosteric drugs (BEN, TH1, TH2, C07, C12) and 20 off-target molecules (STF, GPT, NDA, OXO, CNX, FLQ, PPM, NVB, PA8, P11, P14, ETB, INH, PZA, ETH, STR, DLM, BDQ, LNZ, RIF).

[4] Screening is composed of 6 NSAIDs (CLX, IDT, FBF, IBU, PCT, DCF) and 26 off-target molecules (STF, MFX, GFN, NDA, OXO, FLQ, PPM, LFX, ETP, QPT, IP3, IP1, ZLF, NVB, PA8, P11, P14, ETB, INH, PZA, ETH, STR, DLM, BDQ, LNZ, RIF).

[5] Screening is composed of 6 NSAIDs (CLX, SC5, FBF, IBU, PCT, DCF) and 26 off-target molecules (STP, MFX, GFN, NDA, OXO, FLQ, PPM, LFX, ETP, QPT, IP3, IP1, ZLF, NVB, PA8, P11, P14, ETB, INH, PZA, ETH, STR, DLM, BDQ, LNZ, RIF).

**Table S10. Virtual screening of 3,115 FDA-approved compounds using the *All-at-Once* docking strategy.** The table lists the 148 compounds that emerged as winner across 124 runs, along with their docking success rate based on 100 generated models. Among them, 38 FQs are highlighted in green, while the 8 FQs not selected are also indicated.

**< TableS10-Screening.xls>**

**Table S11. Exploratory application of competitive docking for designing more potent fluoroquinolones (FQs).** The table lists 410 de novo compounds along with their SMILES string and their Tanimoto similarity coefficients relative to sitrafloxacin (STF), sparfloxacin (SPF), fleroxacin (FRX), temafloxacin (TMF), and moxifloxacin (MFX). The success rate of each compound against STF in competitive docking is provided.

**< TableS11-Applying competitive docking to design more potent FQs.xls**

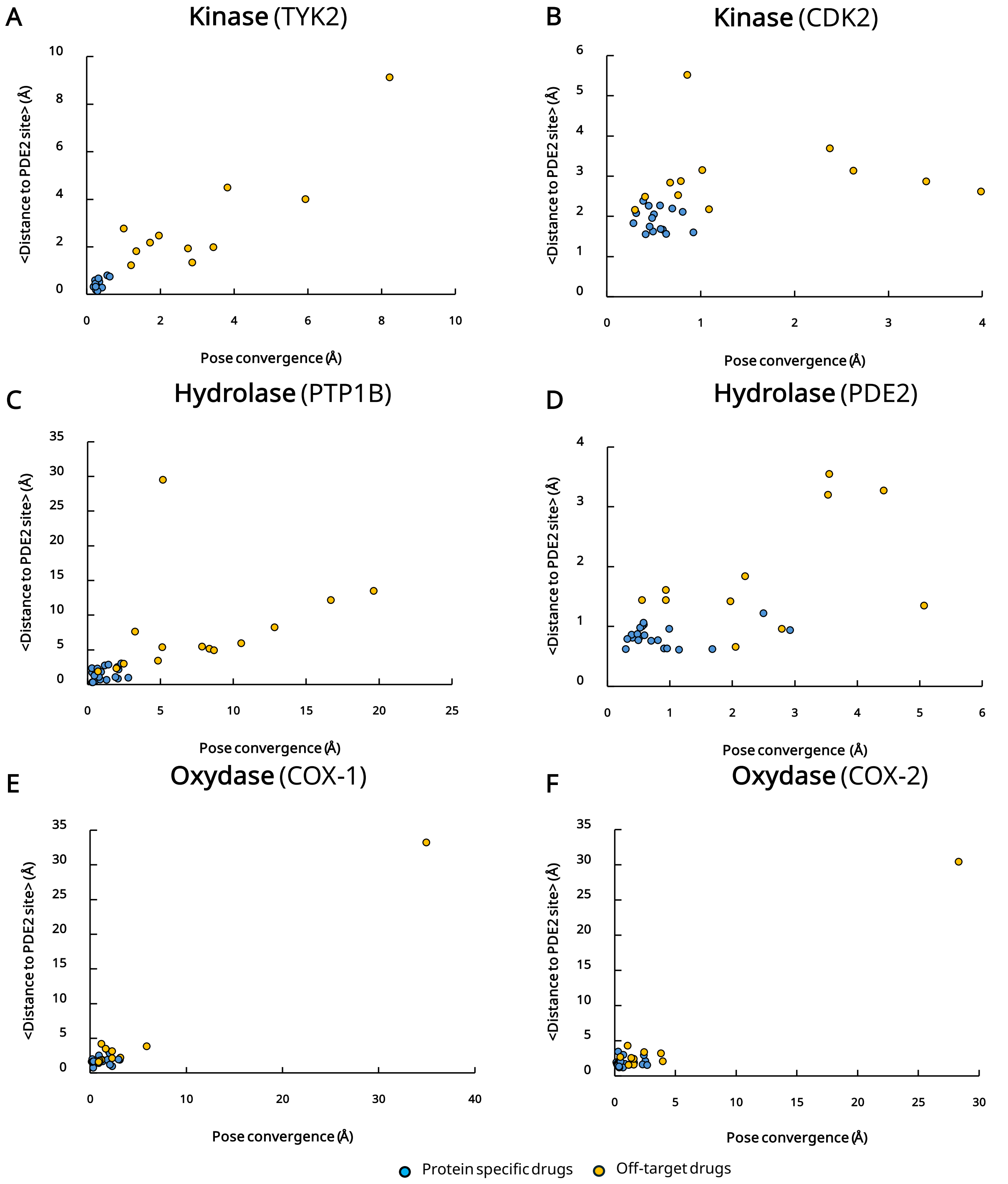


**Fig. S1. Docking specificity of Boltz-2 across six protein benchmarks.** Scatter plots show ligand pose convergence (RMSD) versus binding site distance for true on-target inhibitors (blue) and unrelated off-target compounds (orange). Results are presented for six enzymes: the kinases TYK2 (**A**) and CDK2 (**B**), the hydrolases PTP1B (**C**) and PDE2 (**D**), and the oxidases COX-1 (**E**) and COX-2 (**F**).

| | CNF | CPF | ENX | FLQ | FRX | GFN | GMF | GPF | GRF | LFX | MFX | NDA | NRF | OXO | PFX | PPM | SPF | STF | TMF | TSF | TVF | SCORE |
|---|---|---|---|---|---|---|---|---|---|---|---|---|---|---|---|---|---|---|---|---|---|---|
| CNF | ----- | 22/3 | 23/2 | 22/3 | 6/19 | 5/20 | 24/1 | 24/1 | 22/3 | 22/3 | 8/17 | 25/0 | 25/0 | 24/1 | 25/0 | 23/2 | 5/20 | 4/21 | 3/22 | 2/23 | 2/23 | 316 |
| CPF | 3/22 | ----- | 6/19 | 22/3 | 4/21 | 0/25 | 1/24 | 4/21 | 6/19 | 2/23 | 0/25 | 25/0 | 22/3 | 23/2 | 8/17 | 20/5 | 1/24 | 2/23 | 1/24 | 1/24 | 5/20 | 156 |
| ENX | 2/23 | 19/6 | ----- | 23/2 | 0/25 | 5/20 | 3/22 | 1/24 | 22/3 | 6/19 | 2/23 | 21/4 | 17/8 | 25/0 | 15/10 | 21/4 | 2/23 | 2/23 | 1/24 | 2/23 | 2/23 | 191 |
| FLQ | 3/22 | 3/22 | 2/23 | ----- | 2/23 | 1/24 | 1/24 | 7/18 | 1/24 | 1/24 | 2/23 | 23/2 | 4/21 | 20/5 | 1/24 | 19/6 | 1/24 | 0/25 | 1/24 | 0/25 | 0/25 | 92 |
| FRX | 19/6 | 21/4 | 25/0 | 23/2 | ----- | 19/6 | 23/2 | 23/2 | 24/1 | 25/0 | 2/23 | 25/0 | 23/2 | 25/0 | 24/1 | 22/3 | 0/25 | 3/22 | 7/18 | 20/5 | 17/8 | 370 |
| GFN | 20/5 | 25/0 | 20/5 | 24/1 | 6/19 | ----- | 21/4 | 22/3 | 18/7 | 22/3 | 11/14 | 24/1 | 23/2 | 24/1 | 25/0 | 21/4 | 2/23 | 5/20 | 11/14 | 13/12 | 1/24 | 338 |
| GMF | 1/24 | 24/1 | 22/3 | 24/1 | 2/23 | 4/21 | ----- | 2/23 | 7/18 | 5/20 | 6/19 | 23/2 | 21/4 | 24/1 | 21/4 | 24/1 | 4/21 | 3/22 | 3/22 | 4/21 | 4/21 | 228 |
| GPF | 1/24 | 21/4 | 24/1 | 18/7 | 2/23 | 3/22 | 23/2 | ----- | 4/21 | 9/16 | 2/23 | 24/1 | 21/4 | 20/5 | 22/3 | 23/2 | 3/22 | 0/25 | 2/23 | 3/22 | 2/23 | 227 |
| GRF | 3/22 | 19/6 | 3/22 | 24/1 | 1/24 | 7/18 | 18/7 | 21/4 | ----- | 1/24 | 6/19 | 24/1 | 20/5 | 19/6 | 16/9 | 24/1 | 2/23 | 1/24 | 6/19 | 4/21 | 3/22 | 222 |
| LFX | 3/22 | 23/2 | 19/6 | 24/1 | 0/25 | 3/22 | 20/5 | 16/9 | 24/1 | ----- | 5/20 | 23/2 | 22/3 | 24/1 | 24/1 | 23/2 | 0/25 | 3/22 | 2/23 | 0/25 | 0/25 | 258 |
| MFX | 17/8 | 25/0 | 23/2 | 23/2 | 23/2 | 14/11 | 19/6 | 23/2 | 19/6 | 20/5 | ----- | 22/3 | 23/2 | 25/0 | 23/2 | 21/4 | 18/7 | 6/19 | 24/1 | 21/4 | 24/1 | 413 |
| NDA | 0/25 | 0/25 | 4/21 | 2/23 | 0/25 | 1/24 | 2/23 | 1/24 | 1/24 | 2/23 | 3/22 | ----- | 3/22 | 6/19 | 0/25 | 2/23 | 0/25 | 0/25 | 3/22 | 1/24 | 3/22 | 34 |
| NRF | 0/25 | 3/22 | 8/17 | 21/4 | 2/23 | 2/23 | 4/21 | 4/21 | 5/20 | 3/22 | 2/23 | 22/3 | ----- | 21/4 | 0/25 | 22/3 | 2/23 | 0/25 | 2/23 | 1/24 | 0/25 | 124 |
| OXO | 1/24 | 2/23 | 0/25 | 5/20 | 0/25 | 1/24 | 1/24 | 5/20 | 6/19 | 1/24 | 0/25 | 19/6 | 4/21 | ----- | 1/24 | 1/24 | 1/24 | 0/25 | 2/23 | 0/25 | 2/23 | 52 |
| PFX | 0/25 | 17/8 | 10/15 | 24/1 | 1/24 | 0/25 | 4/21 | 3/22 | 9/16 | 1/24 | 2/23 | 25/0 | 25/0 | 24/1 | ----- | 21/4 | 3/22 | 1/24 | 3/22 | 2/23 | 1/24 | 176 |
| PPM | 2/23 | 5/20 | 4/21 | 6/19 | 3/22 | 4/21 | 1/24 | 2/23 | 1/24 | 2/23 | 4/21 | 23/2 | 3/22 | 24/1 | 4/21 | ----- | 0/25 | 0/25 | 2/23 | 4/21 | 2/23 | 96 |
| SPF | 20/5 | 24/1 | 23/2 | 24/1 | 25/0 | 23/2 | 21/4 | 22/3 | 23/2 | 25/0 | 7/18 | 25/0 | 23/2 | 24/1 | 22/3 | 25/0 | ----- | 3/22 | 6/19 | 23/2 | 18/7 | 406 |
| STF | 21/4 | 23/2 | 23/2 | 25/0 | 22/3 | 20/5 | 22/3 | 25/0 | 24/1 | 22/3 | 19/6 | 25/0 | 25/0 | 25/0 | 24/1 | 25/0 | 22/3 | ----- | 20/5 | 22/3 | 20/5 | 454 |
| TMF | 22/3 | 24/1 | 24/1 | 24/1 | 18/7 | 14/11 | 22/3 | 23/2 | 19/6 | 23/2 | 1/24 | 22/3 | 23/2 | 23/2 | 22/3 | 23/2 | 19/6 | 5/20 | ----- | 18/7 | 19/6 | 388 |
| TSF | 23/2 | 24/1 | 23/2 | 25/0 | 5/20 | 12/13 | 21/4 | 22/3 | 21/4 | 25/0 | 4/21 | 24/1 | 24/1 | 25/0 | 23/2 | 21/4 | 2/23 | 3/22 | 7/18 | ----- | 11/14 | 345 |
| TVF | 23/2 | 20/5 | 23/2 | 25/0 | 8/17 | 24/1 | 21/4 | 23/2 | 22/3 | 25/0 | 1/24 | 22/3 | 25/0 | 23/2 | 24/1 | 23/2 | 7/18 | 5/20 | 6/19 | 14/11 | ----- | 364 |

**Fig. S2. The competition docking matrix of the 21 FQs against *Mtb* DNA gyrase produced with AF3.**

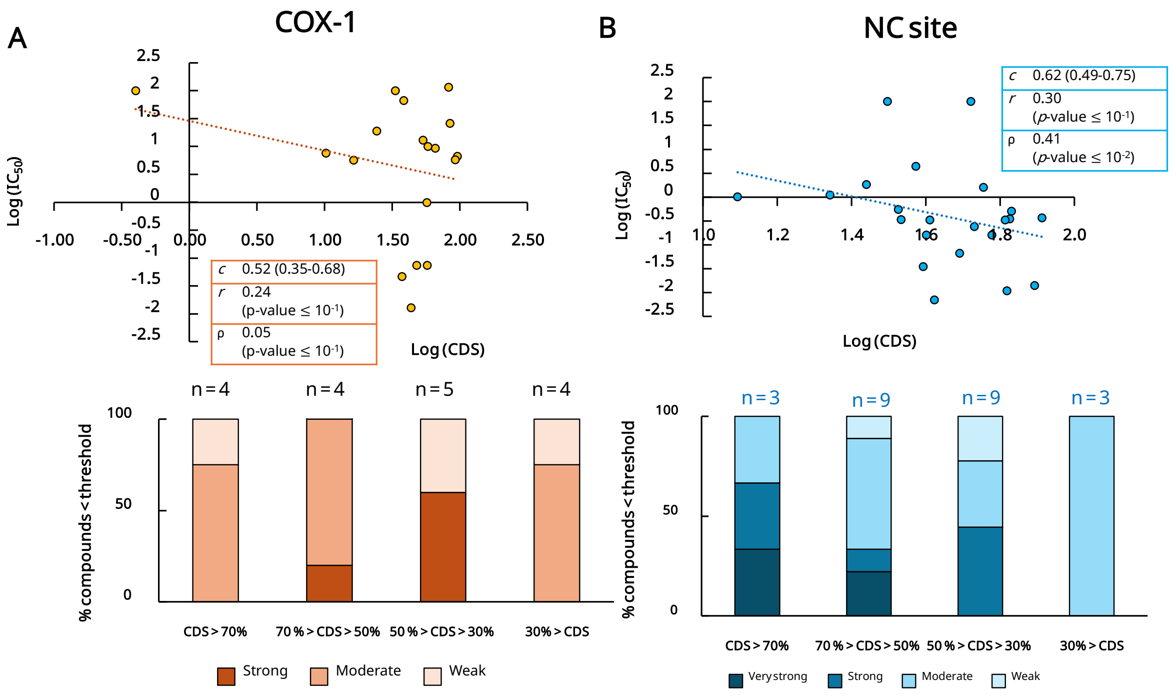


**Fig. S3. Correlation between Competitive Docking Scores (CDS) and experimental inhibitory activities. (A)** NSAIDs against human COX-1. (**B)** NBTIs targeting the NC site of DNA gyrase. The upper panels show scatter plots of CDS versus $IC_{50}$ values, with Pearson's correlation coefficient (*r*), Spearman's rang correlation (ρ) and their respective *p*-values. Rank concordance *c*-index (*c*) with its 95% confidence interval is also reported. An ordinary least squares regression line is included in each scatter plot to visualize the trend. The lower panels show the percentage of NSAIDs and NBTIs within different $IC_{50}$ categories as a function of their CDS ranking ($IC_{50}$ thresholds are listed in Table S5 for NSAIDs and Table S8 for NBTIs. The *n* values indicate the number of inhibitors in each CDS category.

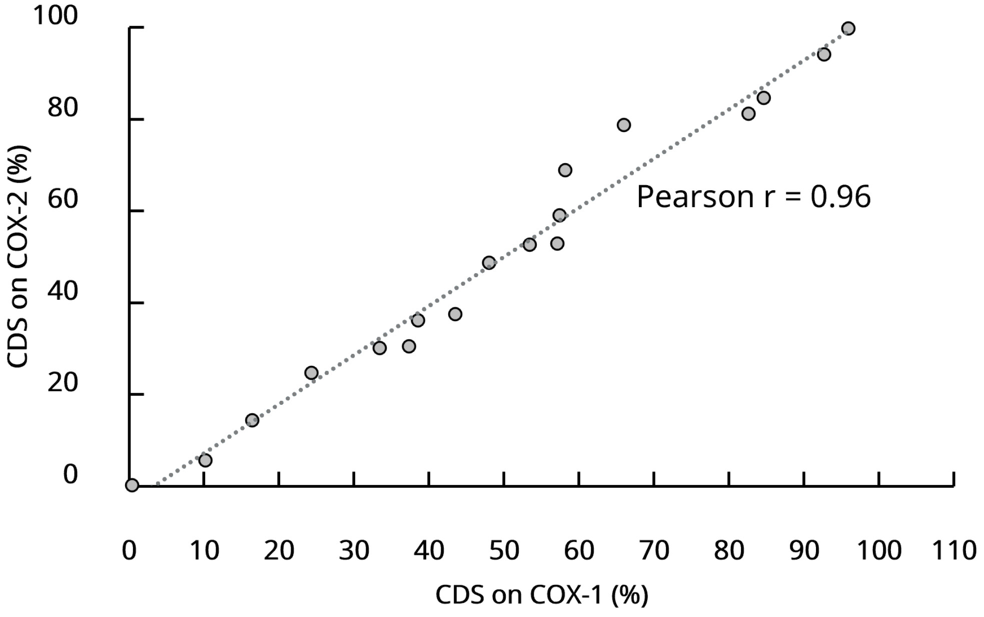


**Fig. S4. Relationship between CDS of inhibitors of COX-1 and COX-2.** The Pearson's correlation coefficient is indicated. CDS were produced with AF3 competitive docking runs.

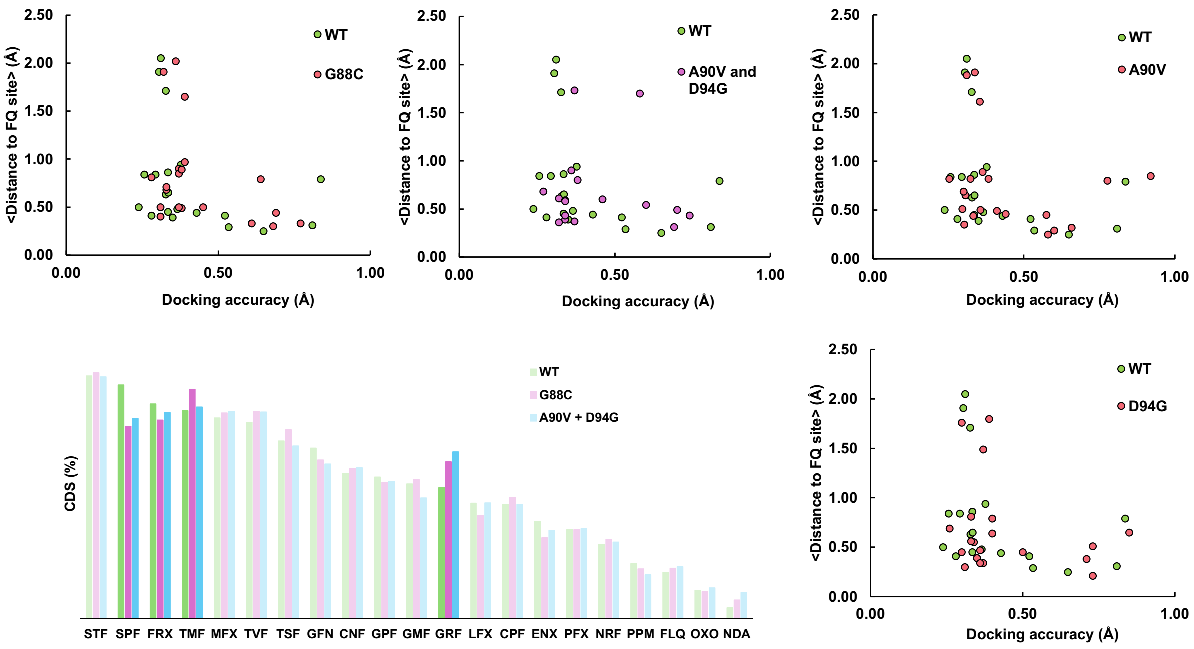


**Fig. S5. Low sensitivity of AF3 docking to point mutations.** Impact of mutations (G88C, A90V, D94G, and A90V+D94G) on docking accuracy and distance from the FQ binding site. FQ positioning shifts occur without a distinct pattern for DNA gyrase variants. Competitive docking scores for G88C and A90V+D94G indicate that rankings are slightly affected by mutations.

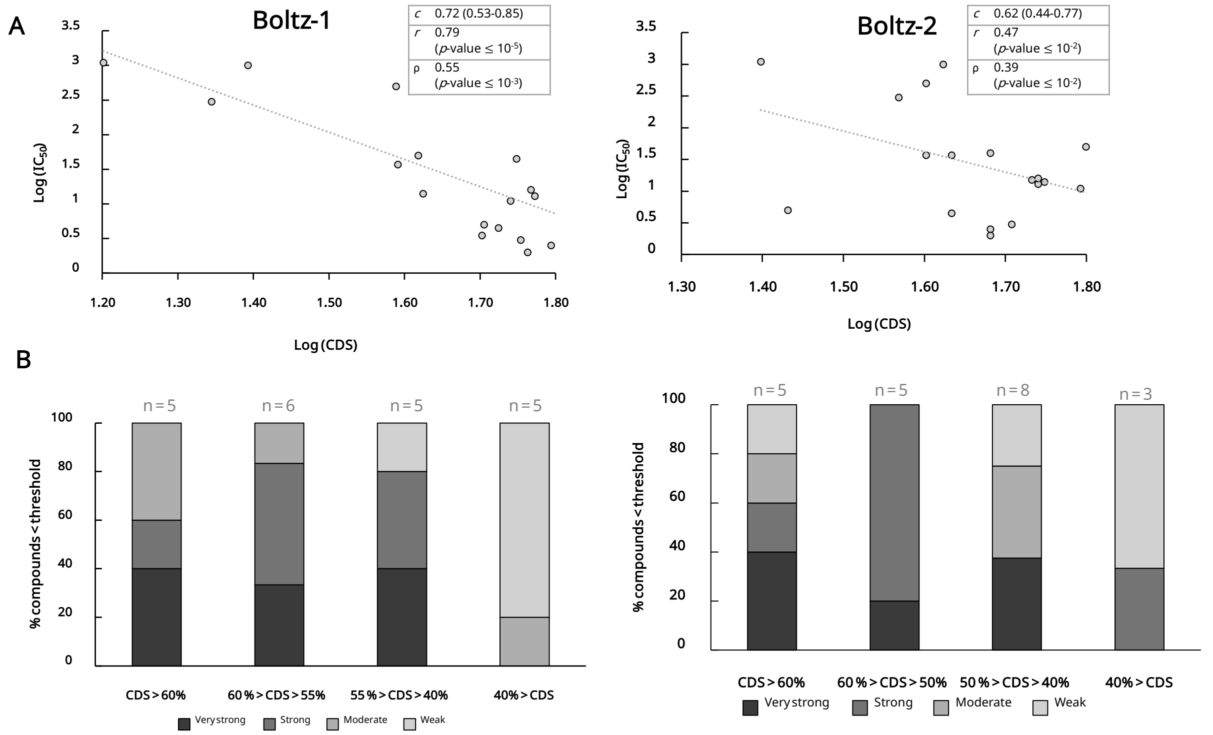


**Fig. S6. Correlation between Competitive Docking Scores (CDS) and experimental inhibitory activities for FQs against *Mtb* DNA gyrase. (A)** Scatter plots illustrate the relationship between the Competitive Docking Score (CDS) and $IC_{50}$ (lighter color) for 21 FQs using Boltz-1 or Boltz-2. The Pearson correlation coefficient ($r$), Spearman's rang correlation ($\rho$) with their corresponding $p$-values, and the rank concordance $c$-index with its 95% confidence interval are reported. An ordinary least squares regression line is included in each scatter plot to visualize the trend. (**B**) Percentage of FQs within $IC_{50}$ threshold categories, stratified by CDS ranking using Boltz-1 or Boltz-2. The bar chart shows the percentage of FQs falling into four $IC_{50}$ threshold categories, based on their CDS ranking. $IC_{50}$ thresholds are detailed in Table S3. The 'n' values indicate the number of FQs in each CDS category.

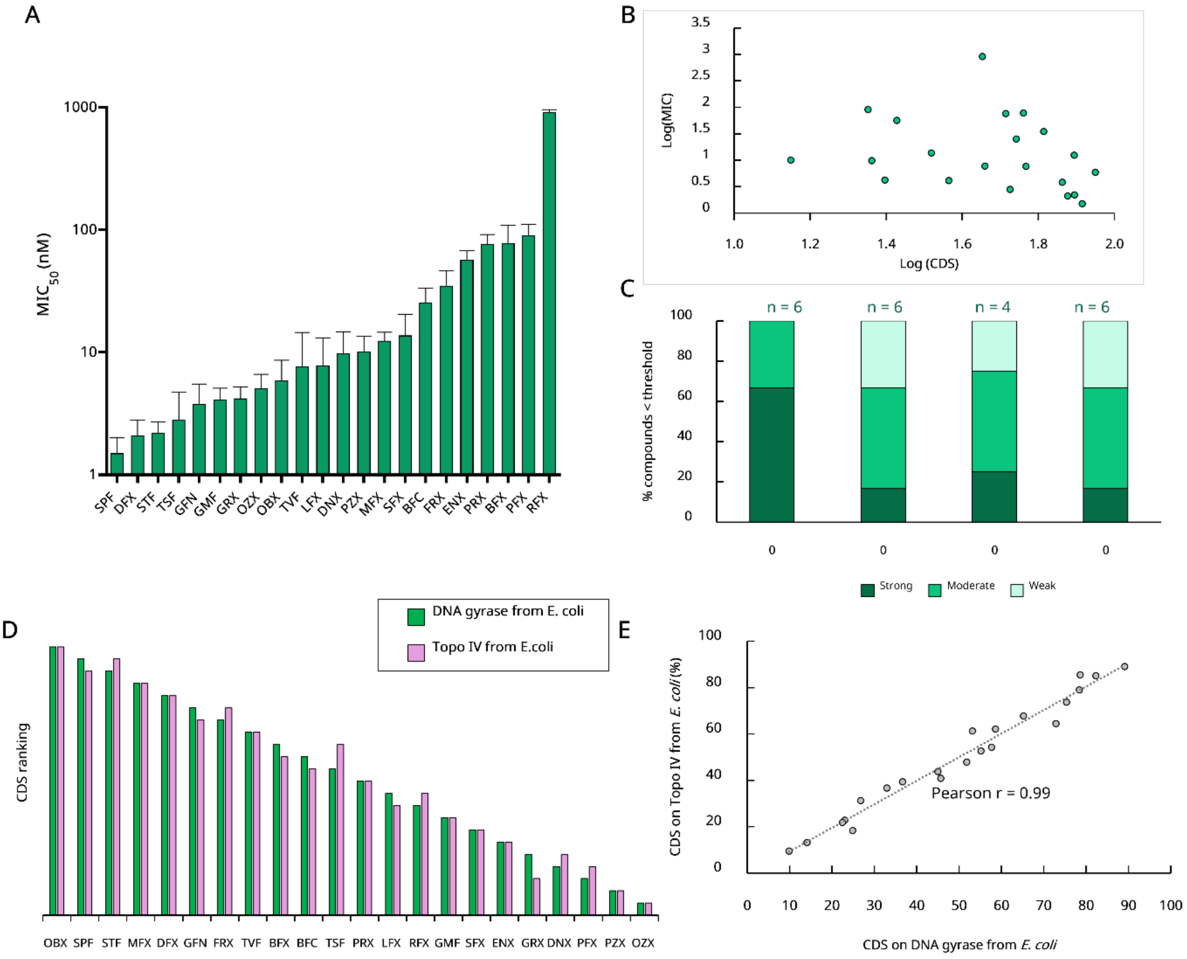


**Fig. S7. Experimental inhibitory activity of FQs against *E. coli* and correlation with CDS rankings.** (**A**) Bar graph displaying the measured $MIC_{50}$ values (in nM, log scale) for 22 FQs against *E. coli*. Values represent the mean of at least three biological replicates, with standard deviation indicated. (**B**) Scatter plot illustrating the correlation between Competitive Docking Score (CDS) and $MIC_{50}$ values for the 22 FQs using AF3. (**C**) Distribution of FQs across $MIC_{50}$ threshold categories, stratified by CDS ranking using AF3. $MIC_{50}$ thresholds are defined in Table S4. 'n' values denote the number of FQs in each CDS category. CDS ranking comparison between *E. coli* DNA gyrase and topoisomerase IV produced by AF3 with (**D**) bar graph of both CDS rankings and (**E**) Scatter plot of CDSs.

**A**

| Drug | Tanimoto from STF | % Win vs STF | Synthetic accessibility | Solubility | Druglikeness | |
|---|---|---|---|---|---|---|
| | | | | | Lipinski | Ghose |
| Z03 | 0.31 | 86.0 | 3.81 | Soluble | Yes | Yes |
| Z06 | 0.26 | 76.0 | 4.11 | Soluble | Yes | Yes |
| Z36 | 0.25 | 80.0 | 3.75 | Moderate | Yes | Yes |
| Z46 | 0.35 | 78.0 | 3.72 | Moderate | Yes | Yes |
| Z53 | 0.26 | 76.0 | 4.00 | Soluble | Yes | Yes |
| Z99 | 0.44 | 70.0 | 3.96 | Soluble | Yes | Yes |
| Z111 | 0.88 | 74.0 | 4.44 | Moderate | Yes | Yes |
| Z112 | 0.88 | 82.0 | 4.41 | Moderate | Yes | Yes |
| Z119 | 0.88 | 70.0 | 4.44 | Moderate | Yes | Yes |
| Z121 | 0.30 | 70.0 | 3.72 | Soluble | Yes | Yes |
| Z128 | 0.33 | 88.0 | 2.99 | Moderate | Yes | Yes |
| Z181 | 0.36 | 82.0 | 3.77 | Soluble | Yes | Yes |
| Z184 | 0.32 | 82.0 | 3.85 | Soluble | Yes | Yes |
| Z187 | 0.35 | 78.0 | 3.62 | Moderate | Yes | Yes |
| Z188 | 0.35 | 74.0 | 3.62 | Moderate | Yes | Yes |
| Z196 | 0.35 | 70.0 | 3.92 | Moderate | Yes | Yes |
| Z226 | 0.35 | 84.0 | 4.13 | Soluble | Yes | Yes |
| Z242 | 0.27 | 82.0 | 3.78 | Soluble | Yes | Yes |
| Z246 | 0.81 | 86.0 | 4.62 | Soluble | Yes | Yes |
| Z247 | 0.35 | 92.0 | 4.47 | Soluble | Yes | Yes |
| Z248 | 0.35 | 86.0 | 4.04 | Soluble | Yes | Yes |
| Z249 | 0.38 | 88.0 | 4.15 | Soluble | Yes | Yes |
| Z282 | 0.27 | 74.0 | 3.73 | Moderate | Yes | Yes |
| Z305 | 0.27 | 74.0 | 4.05 | Soluble | Yes | Yes |
| Z306 | 0.27 | 84.0 | 3.79 | Moderate | Yes | Yes |
| Z336 | 0.32 | 80.0 | 4.09 | Moderate | Yes | Yes |
| Z365 | 0.26 | 90.0 | 4.02 | Moderate | Yes | Yes |
| Z366 | 0.33 | 84.0 | 3.94 | Soluble | Yes | Yes |
| Z375 | 0.26 | 72.0 | 3.81 | Soluble | Yes | Yes |
| Z384 | 0.76 | 74.0 | 4.36 | Moderate | Yes | Yes |
| Z396 | 0.26 | 82.0 | 4.05 | Soluble | Yes | Yes |

**B**

The chemical structures are not shown at this stage due to pending technology disclosure. They will be made available upon publication of the article

**Fig. S8.** ***In silico*** **design of more efficient compounds. (A)** Compounds generated by applying a >70% CDS score against the reference STF compound. ADMET profiling of candidate compounds using predictive filters implemented in the SwissADME tool[19]. Synthetic accessibility is scored on a scale from 1 (easy to synthesize) to 10 (difficult), while solubility class is determined using the SILICOS-IT. **(B)** Structure of the heigh molecules passing the solubility filter and the synthetic accessibility (< 4.0). The acronyms used in this study are indicated below each structure. Chemical structures were drawn and visualized using Marvin (Marvin 24.3.2, ChemAxon, https://www.chemaxon.com).
